# Time-resolved lineage recording reveals a pre-existing, heritable cell state underlying metastatic potential

**DOI:** 10.64898/2026.08.10.744013

**Authors:** Jihye Park, Yoojin Chang, Joshua Schiffman, Divya Koyyalagunta, Hrishikesh Somayaji, Caden N. McQuillen, Joseph Chan, Quaid Morris, Dan A. Landau, Hyongbum Henry Kim, Junhong Choi

**Affiliations:** Developmental Biology Program, Memorial Sloan Kettering Cancer Center, New York, United States; Department of Pharmacology, Yonsei University College of Medicine, Seoul 03722, South Korea; New York Genome Center, New York, NY, USA; Sandra and Edward Meyer Cancer Center, Weill Cornell Medicine, New York, NY, USA; Computational and Systems Biology Program, Memorial Sloan Kettering Cancer Center, New York, United States; Human Oncology and Pathogenesis Program, Memorial Sloan Kettering Cancer Center, New York, United States; Graduate School of Medical Science, Brain Korea 21 Project, Yonsei University College of Medicine, Seoul 03722, South Korea; Severance Biomedical Science Institute, Yonsei University College of Medicine, Seoul 03722, South Korea; Center for Nanomedicine, Institute for Basic Science (IBS), Seoul 03722, South Korea; Yonsei-IBS Institute, Yonsei University, Seoul 03722, South Korea; Institute for Immunology and Immunological Diseases, Yonsei University College of Medicine, Seoul 03722, South Korea; Woo Choo Lee Institute for Precision Drug Development, Yonsei University College of Medicine, Seoul 03722, South Korea; Won-Sang Lee Institute for Hearing Loss, Yonsei University College of Medicine, Seoul 03722, South Korea

**Author notes:** These authors contributed equally (Jihye Park, Yoojin Chang). Correspondence (H.H.K.), (J.Choi).

## Abstract

Metastasis causes most cancer deaths^1,2^, yet no recurrent mutation specifically drives it^3,4^, raising the possibility that metastatic potential is a non-genetic yet heritable cell state. Classic experiments established that metastatically predisposed subclones pre-exist within a tumor and that these predispositions are inherited over many cell divisions^5^, but what molecular states or factors underlie this predisposition remain unknown. While previous lineage recording studies^6,7^ mapped how tumors disseminate, their recording sites saturate too quickly to resolve when a lineage branched, or to attribute a state to its founder. Here we show, using a DNA Typewriter lineage recorder^8^ with nearly 1,000 recording sites in lung cancer cells, that metastatic potential is already present before dissemination, with colonization predicted by a pre-existing glycolytic state and further spread by expression of *ENO1*, a glycolytic enzyme that also moonlights as a cell-surface plasminogen receptor^9^. Profiling the pre-transplant cells and the post-transplantation tumors for both their transcriptomes and their lineage recordings, we reconstructed time-resolved lineage trees across three orthotopically transplanted mice. These trees trace each liver metastasis to a single founder of known pre-transplant state, dating each dissemination event from the primary lung. When every clone was scored before transplant against 349 genes recurrently heritable in vitro, both that set and the glycolytic state shifted a clone’s odds of colonizing the lung, each holding when the two were fitted together. At the gene level, sixteen genes were both heritable and predictive of colonization, and *ENO1* alone also predicted which established clones spread further. Hypoxia, the program most strongly associated with phylogenetic fitness within the metastases, did not predict colonization when scored before transplant, separating niche-selected traits from the inherited cell state. Metastatic potential in this system is therefore transmitted along the lineage rather than acquired after seeding. Looking forward, we anticipate that time-resolved lineage recorders will enable the separation of the heritable and acquired components of the cellular heterogeneity seen in single-cell studies of tumor progression and drug tolerance.

## INTRODUCTION

Cancer metastasis is responsible for the majority of cancer-associated deaths^1,2^, yet the events that produce a metastasis are hardly trackable. A founding cell (or cells) leaves the primary tumor, survives transit, and re-establishes growth at a distant site. By the time a metastasis is detected, the cellular traits that drove it are gone. In a classic experiment, Fidler and Kripke^5^ combined a Luria-Delbrück fluctuation analysis^10^ with serial subcloning to establish two things: that a tumor is a heterogeneous population of cells with varying metastatic potential, and that this potential is heritable across many generations in culture. What has remained open ever since is the basis of that heritable trait. Despite extensive genomic characterization of primary and metastatic tumors, no recurrent mutations that specifically drive the metastatic cascade have been identified^3,4^, pointing to a non-genetic basis, but which molecular state or cellular program encodes it has been hard to establish, because it requires reconstructing the state of the founding cells that gave rise to a metastasis after the fact.

In addition to a founder’s state, the history of a metastasis is similarly lost through the endpoint sampling of a metastatic tumor. A tumor can seed early or late, through single or multiple migration events, as single cells or multicellular clusters, and by direct or stepwise routes with varying organotropism^11,12^. In humans, this history can be reconstructed retrospectively from the mutations a tumor happens to carry: phylogenetic analyses of paired primary tumors and metastases show that lymph node metastases arise through a wider evolutionary bottleneck than distant metastases^13,14^ and that distant lesions tend to seed late and largely directly from the primary^15^. Their resolution is set by mutational diversity, however, so dating individual events is difficult, and they cannot connect a metastasis to the state of the cell that founded it. Recovering both the timing of dissemination and the state of the founding cells therefore requires a high-resolution record of cell-lineage relationships written into the cells themselves.

Engineering cancer cells with lineage recorders addresses this by writing heritable marks into the genome that accumulate as cells divide, so that the genealogical relationships between primary and metastatic tumor cells can be reconstructed from the marks alone. Applied to xenograft metastasis, CRISPR-Cas9 recorders have mapped the rates and routes of dissemination in lung adenocarcinoma^6^ and identified selection for hybrid epithelial-mesenchymal states^7^. Cas9 scarring is nonetheless unordered and saturates a small number of target sites quickly, so it records topology but not the time at which branches formed. And neither study could establish whether it was set before dissemination: one induced recording only after transplantation^7^, while the other, though it began recording before implantation and examined pro-metastatic genes in the pre-implantation pool^6^, lacked the lineage resolution to show that this expression was heritable rather than transient.

Prime editing-based lineage recorders, such as DNA Typewriter^8^ and PEtracer^16^, address the first of these limitations by inserting predefined barcodes at multiple genomic sites that can be read as synthetic mutations to reconstruct lineage relationships among cells. DNA Typewriter, the system we use here, arranges these sites as a tandem array and writes them directionally, so each site is filled only after the preceding one. Because the insertions are ordered and accumulate steadily, their depth along the array reports how much time has elapsed, and once the editing rate is calibrated the number of accumulated edits at a lineage branch converts to absolute time since the onset of the recording. Realizing this for cancer metastasis requires a recording line with enough capacity to keep editing for the full duration of an in vivo experiment without early saturation, and a way to read the recorder together with the transcriptome of the same cell.

To follow metastasis of orthotopically transplanted lung cancer cells, here we engineered a high-capacity DNA Typewriter recorder in NCI-H1299 cells with 996 recording sites (166 arrays of 6 sites each) that continues to edit for 45 days, including the 5-week in vivo window, and marked each founding clone with a static clonal barcode. At the time of transplantation, we profiled around 40% of the recording pool by single-cell RNA sequencing, which gives every clone a transcriptional state measured on the day of transplant, and continued to culture the remaining cells to measure editing rate throughout the experiment and to identify recurrently heritable across clonal generations. At the end of the in vivo experiment, we harvested tumors from three mice and recovered lineage recordings and transcriptome from the same cells. Reconstructed time-resolved lineage trees dated individual metastatic disseminations and show that metastatic competence in this model is a pre-existing, heritable cell state present in the founding cells before they leave the primary tumor. A glycolytic state and an aggregate score over 349 recurrently heritable genes each shift the odds that a clone colonizes the lung, while *ENO1* alone predicts which established clones go on to spread. Hypoxia, the program most strongly associated with phylogenetic fitness within the metastases, did not predict colonization when scored before transplant, which separates a trait selected within the niche from the state inherited along the lineage.

## RESULTS

### Engineering an NCI-H1299 line with a high-capacity DNA Typewriter lineage recorder

As a tractable model for lung cancer metastasis, we selected NCI-H1299, a non-small cell lung carcinoma (NSCLC) line with loss of p53 expression and an activating NRAS Q61K mutation. These cells display epithelial-like morphology, proliferate rapidly (doubling time approximately 22 to 30 hours^17,18^), and form metastases in immunodeficient hosts, making them a well-characterized xenograft model.

We engineered these cells for DNA Typewriter recording by integrating a Cre-activatable Prime Editor (LSL-PEmax-P2A-mClover3) together with pegRNA cassettes encoding an array 6 sites (TAPE) using the piggyBAC transposon system (**Supplementary Table S1**), and screened monoclonal candidates for TAPE copy number and Prime Editor activation (**Supplementary Figure 1a,b**; **Methods**). The selected founder line, Founder-39, carried 166 independent TAPE arrays of up to six ordered sites each (996 recording sites in total) and accumulated an average of around 254 insertions per cell over 7 days of editing, with fewer than 1.4% of 6-site arrays saturating, providing a monoclonal line with the recording capacity to track metastasis in vivo.

### Tracking lung cancer metastasis with combined clonal and lineage barcoding

Recording from a single founding cell would in principle capture every subsequent division and reconstruct a complete genealogy from one common ancestor, but this is impractical for xenografts, which need tens of thousands of cells per injection: expanding a single cell to that number would take two to three weeks and consume recording capacity needed in vivo. We therefore began from a pool of founder clones, each marked with a distinct static clonal barcode (ClonalBC) introduced by low-MOI lentiviral transduction (**Figure 1a; Supplementary Table S2**); the ClonalBC construct also encoded luciferase for IVIS bioluminescence imaging and a puromycin-resistance cassette (**Supplementary Figure 1c**). The static ClonalBC reports which founding clone a cell belongs to, while the evolving TAPE resolves the subclonal genealogy within that clone.

**Figure 1.**
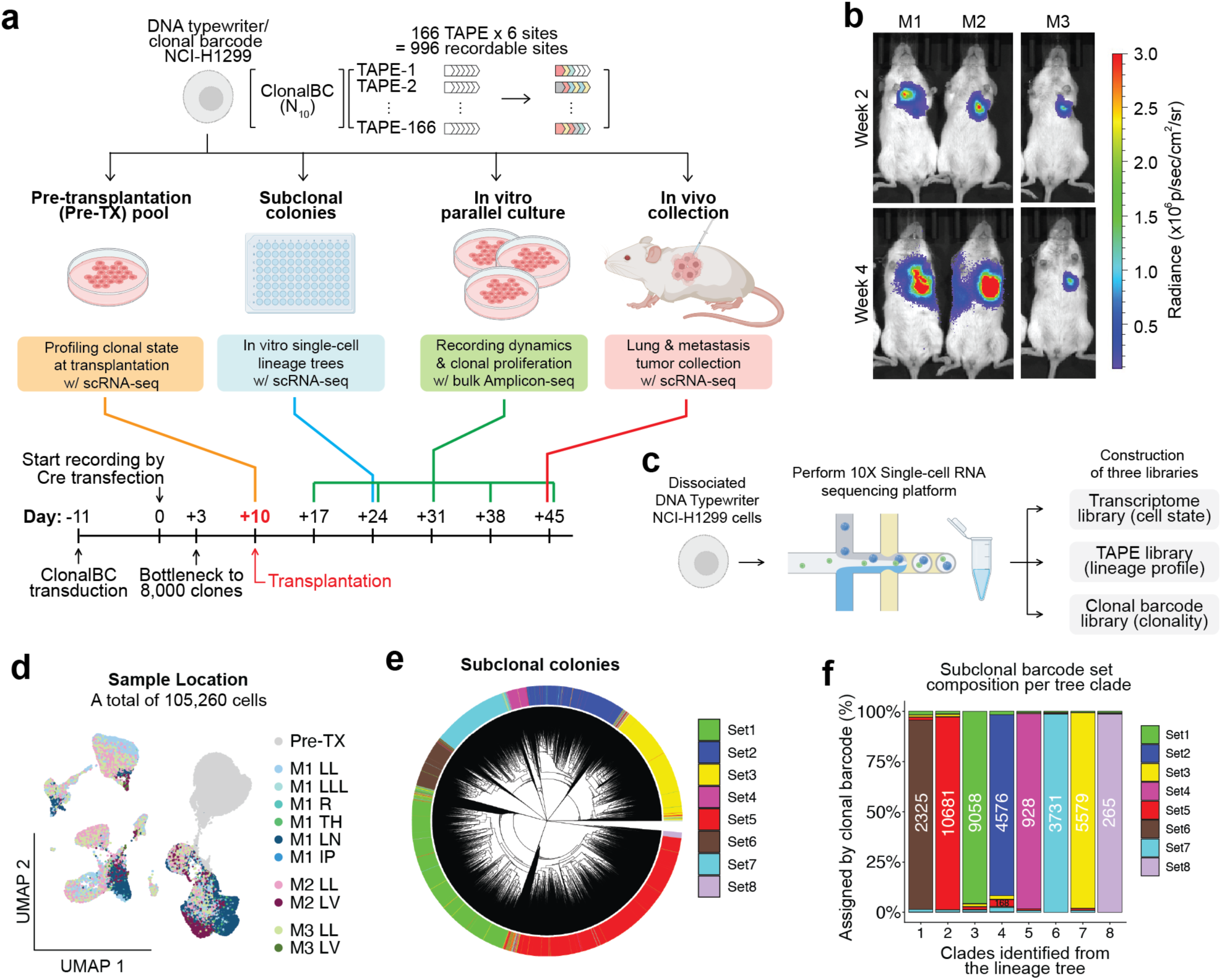
Lineage recording during lung cancer metastasis. (a) Experimental design and timeline. Founder-39 NCI-H1299 cells, carrying 166 TAPE arrays (996 recordable sites) and a static clonal barcode (ClonalBC), were split into four branches: orthotopic transplantation into the left lung of three mice (In vivo transplantation), scRNA-seq profiling at transplantation (Pre-transplantation pool, Pre-TX), a continued parallel in vitro culture with weekly bulk amplicon-seq (In vitro parallel culture), and single-cell-derived cultures for in vitro lineage-tree reconstruction (Subclonal colonies). Timeline, with day 0 the initiation of recording by Cre transfection: ClonalBC lentiviral transduction (day –11), Cre activation (day 0), bottlenecking to ∼8,000 clones (day 3), orthotopic transplantation (day 10), and harvest (day 45, 35 days after transplantation). (b) Representative IVIS bioluminescence images of the three mice (M1 to M3) at weeks 2 and 4 after transplantation. (c) Single-cell processing. Dissociated, FACS-purified mRFP-positive tumor cells were profiled on the 10x Genomics platform, and three libraries were prepared from the resulting cDNA: the transcriptome (cell state), TAPE insertions (lineage), and the clonal barcode (clonality). (d) UMAP of the 105,260 single cells profiled across all samples, colored by sample of origin (Pre-TX and each mouse and organ). (e) Lineage tree of the eight recovered subclonal colonies reconstructed from TAPE insertions alone, with the outer ring coloring each cell by its clonal-barcode set (sets 1 to 8). Ten wells were expanded and profiled; two recovered only 14 and 22 cells and are not shown. (f) Subclonal barcode-set composition of each clade from the tree in (e), with the number of cells per clade above each bar. Each clade is composed almost entirely of a single well’s barcode set, so clades defined by the recorder correspond to the independently introduced clonal barcodes.

**Supplementary Figure 1.**
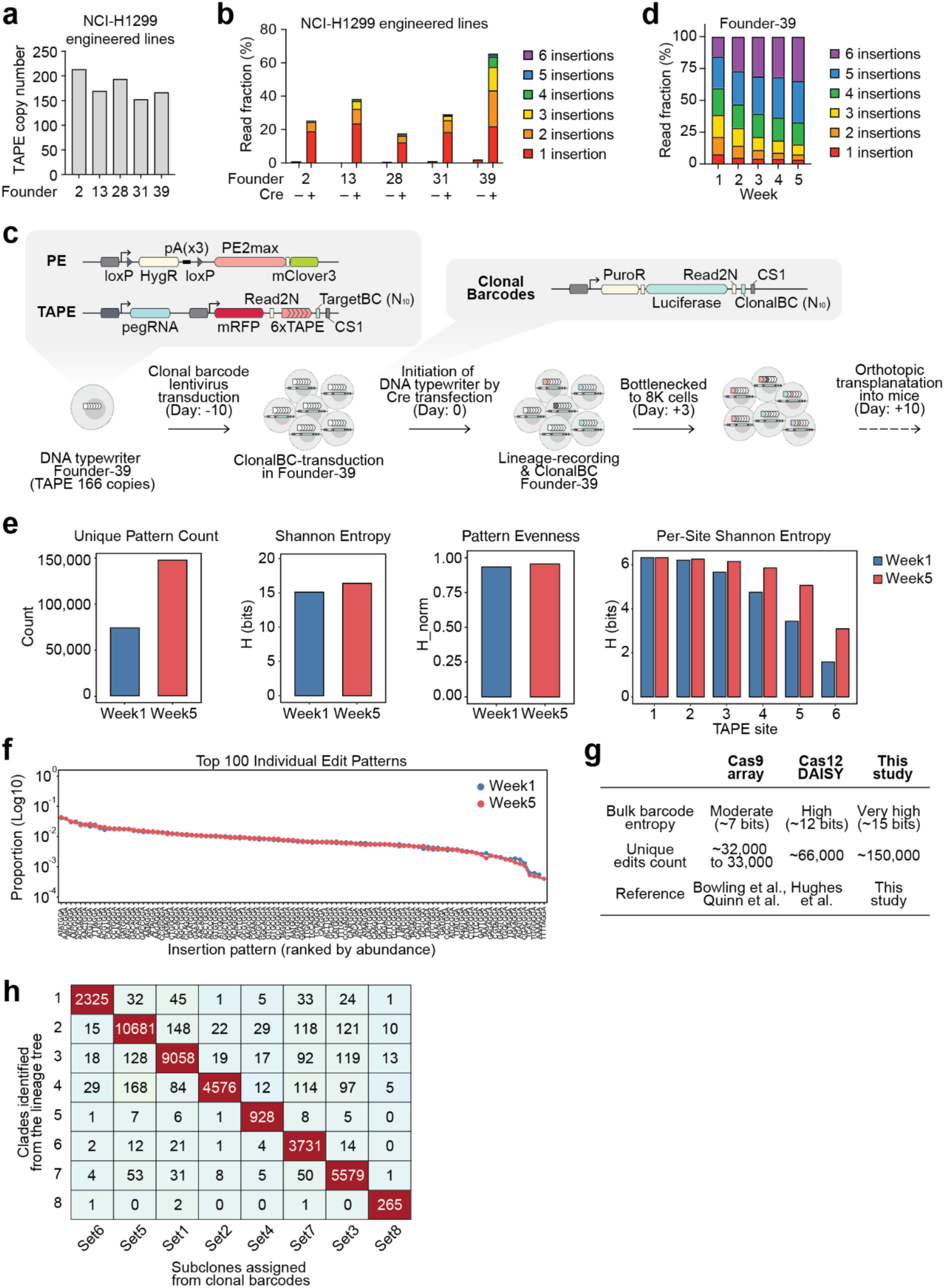
Engineering and benchmarking of the DNA Typewriter recording line. (a) Estimated TAPE copy number in each candidate NCI-H1299 line (five top ones among 48 screened shown), from the number of unique TargetBCs recovered by bulk amplicon sequencing. (b) Recording efficiency and specificity in each candidate line without and with Cre transfection (Cre −/+), scored as the read fraction with 1 to 6 insertions. (c) Construct schematics and workflow for generating clonally barcoded recording cells: the Cre-activatable Prime Editor (LSL-PEmax-P2A-mClover3), the 6xTAPE recorder (pegRNA-mRFP-6xTAPE-TargetBC), and the clonal-barcode construct (luciferase-ClonalBC-puromycin resistance), with the timeline (ClonalBC transduction day –11, Cre activation day 0, bottlenecking day 3, transplantation day 10). (d) Recording by Founder-39 over the five-week window matching the in vivo experiment (day 17 to day 45), as the read fraction with 1 to 6 insertions per week. (e) Recording diversity of Founder-39 at day 17 versus day 45 (weeks 1 and 5): unique pattern count, Shannon entropy, pattern evenness, and per-site Shannon entropy. Unique patterns increase and editing continues to reach the later TAPE sites while evenness stays high. (f) Frequencies of the individual insertion sequences (NNNNGGA) at day 17 versus day 45, showing a stable insertion spectrum over time. (g) Comparison of DNA Typewriter recording with Cas9-array and Cas12a (DAISY) lineage recorders by barcode size, bulk barcode entropy, and number of unique edits. (h) Cross-tabulation of the subclone lineage-tree clades (rows) against the clonal-barcode sets (columns), with the number of cells in each clade-barcode pair. The near-diagonal structure shows that each clade corresponds to a single clonal-barcode set.

**Supplementary Figure 2.**
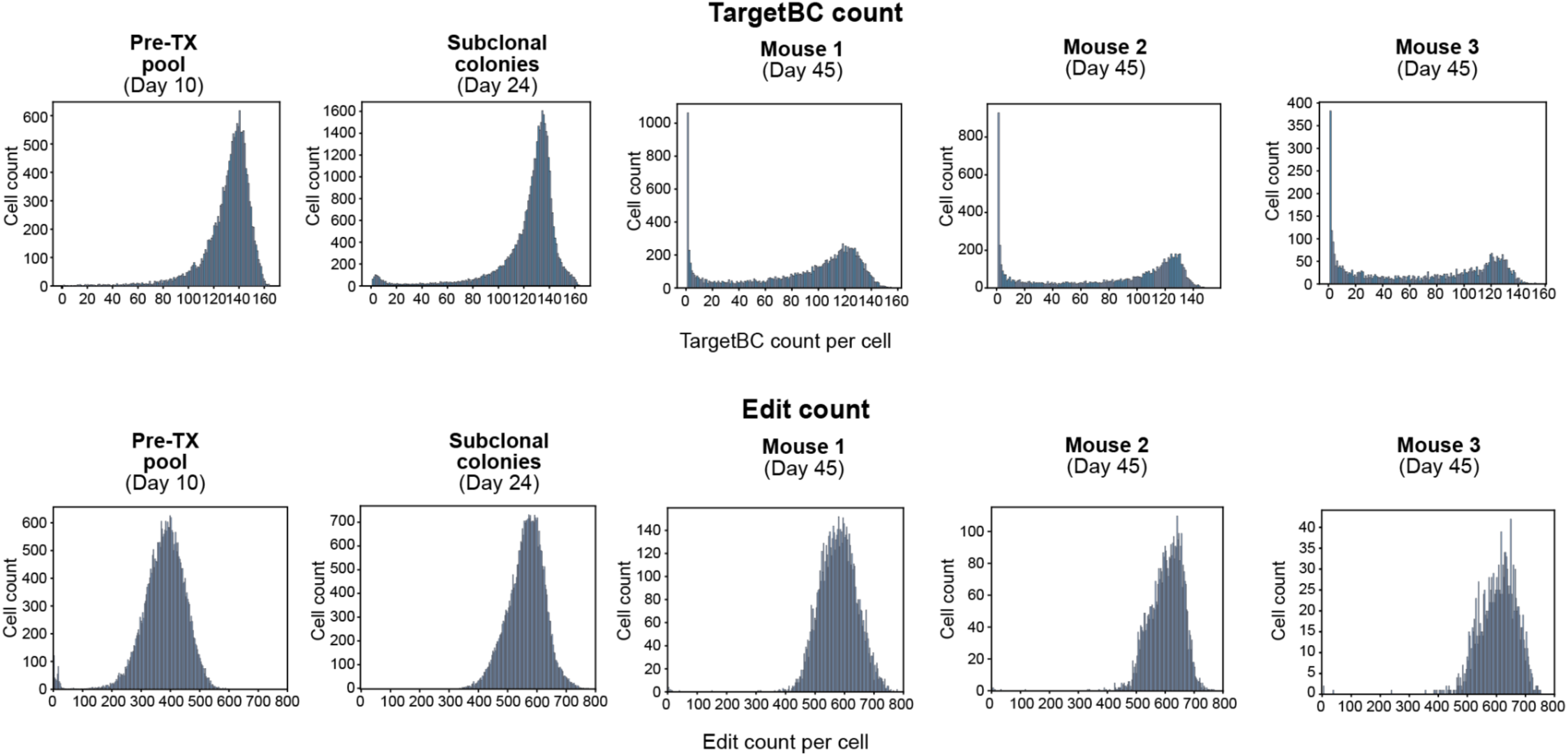
Per-cell recovery of lineage barcodes across samples. For each sample (the pre-transplantation pool, the in vitro subclonal colonies, and Mouse 1 to Mouse 3), distributions of (top) the number of TargetBCs recovered per cell, and (bottom) the number of TAPE insertions recovered per cell. In vitro cells recover most of the 166 TAPEs and several hundred edits per cell, and edit counts rise from the early pool to the week-5 in vivo samples.

We set day 0 as the initiation of recording by Cre transfection, transplanted at day 10, and harvested at day 45 (35 days after transplantation; **Figure 1a**). After bottlenecking the barcoded pool to ∼8,000 cells and expanding it ∼20-fold to ∼160,000 so that multiple siblings of each clone were represented, we split it into four branches: 5,000–10,000 cells were orthotopically injected into the left lung of each of three mice (Mouse 1 to Mouse 3); 40% was profiled by scRNA-seq at transplantation to capture pre-injection states (Pre-transplantation pool, Pre-TX); a small aliquot was single-cell-sorted to expand subclone cultures for in vitro lineage-tree reconstruction (Subclonal colonies); and the remainder was maintained as a parallel in vitro culture over five weeks with weekly barcode profiling (In vitro parallel culture). Tumor growth in each mouse was tracked by biweekly IVIS bioluminescence imaging (**Figure 1b**). At day 45, we imaged each mouse for metastatic foci, isolated mRFP-positive Founder-39 cells from each organ, and profiled them by scRNA-seq (10x Genomics), reading out the transcriptome, ClonalBC, and TAPE insertions as three separate sequencing libraries (**Figure 1c**) for 105,260 single cells across all samples (**Figure 1d**).

### Validating sustained recording and clonal-lineage concordance

Across every sample type (the Pre-TX pool, the single-cell subclones, and the three mice), targeted enrichment recovered abundant TargetBCs and TAPE insertions per cell with high number of accumulated edits (**Supplementary Figure 2**), confirming that lineage recordings were captured efficiently from the scRNA-seq data. The in vitro parallel branch let us test recording over the same five-week window as the in vivo experiment: insertions accumulated each week without saturating (**Supplementary Figure 1d**), and recording diversity grew rather than plateaued, with unique insertion patterns roughly doubling from week 1 to week 5 (∼75,000 to ∼150,000) while the insertion spectrum itself remained stable (**Supplementary Figure 1e,f**). Benchmarked against other CRISPR-based recorders, Founder-39 reached ∼15 bits of barcode entropy and ∼150,000 unique edit patterns, roughly double Cas12a DAISY and several-fold above Cas9 arrays^6,19–21^ (**Supplementary Figure 1g**).

Finally, we asked whether the two barcode modalities report concordant lineages. Ten subclone wells were expanded and profiled, of which eight yielded enough cells for analysis; the remaining two recovered only 14 and 22 cells. Pooling the eight, we reconstructed a lineage tree from the TAPE insertions alone, without reference to the clonal barcodes (**Figure 1e**). Each well resolved as a distinct clade composed almost entirely of cells carrying a single clonal-barcode set (**Figure 1f**; **Supplementary Figure 1h**). Because the topology was inferred from the recorder alone, this correspondence validates the resulting lineage structure.

### Metastatic dissemination varies widely and is uncoupled from proliferation rate

Having validated the two barcodes as concordant (**Figure 1e,f**), we applied the system to the three xenografted mice, surveying the primary lung and the organs that commonly receive metastases at harvest (lymph nodes, thoracic mass, lower left lung, right lung, intraperitoneal nodes, and liver; **Figure 2a**). Five weeks after transplantation, mRFP fluorescence marked the primary tumor in the left lung and metastatic cells across these organs, and the three mice differed markedly in extent (**Figure 2b**). Mouse 1 disseminated broadly across the thoracic cavity, concentrating in the mediastinal and regional lymph nodes and reaching across the thoracic mass, intraperitoneal nodes, the right lung, and the lower left lung. Metastasis in Mouse 2 and Mouse 3 were instead focal, with defined colonies confined to the primary lung and the liver and only a handful of clonal barcodes recovered at the metastatic sites (**Figure 2b,c**).

**Figure 2.**
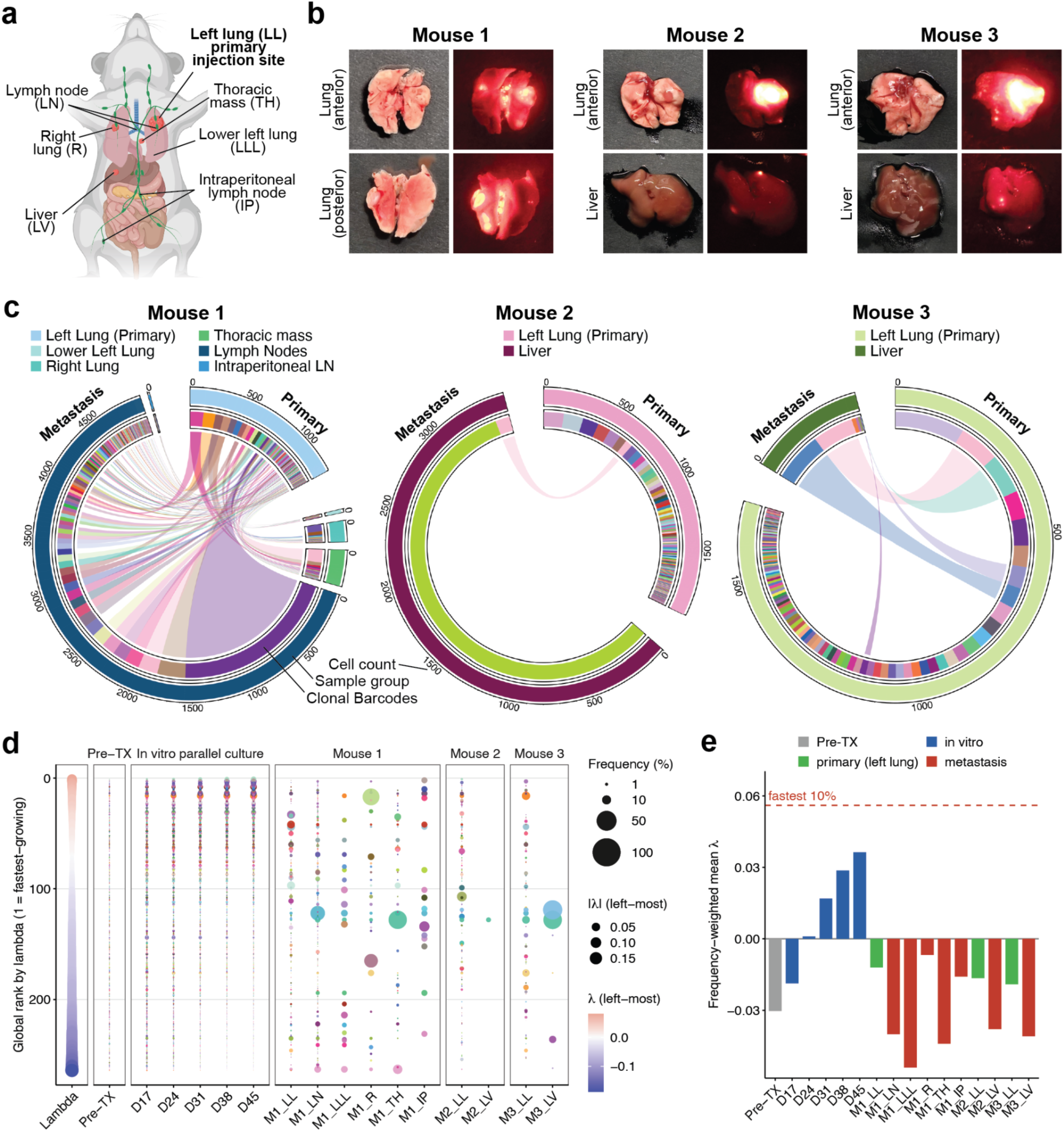
Metastatic dissemination varies widely in extent and is not selected by clone proliferation rate. (a) Anatomical schematic of the transplant and harvest sites: the left lung (LL) primary injection site and the organs surveyed for metastasis, including the left lung lower (LLL), right lung (R), lymph nodes (LN), thoracic mass (TH), intraperitoneal lymph node (IP), and liver (LV). (b) Representative paired brightfield and mRFP-fluorescence images of the primary lung and metastatic organs for each mouse (Mouse 1, lung anterior and posterior; Mouse 2 and Mouse 3, lung and liver), showing the mRFP-positive primary tumor and fluorescent metastatic foci. (c) Clonal composition of the primary and metastatic cells from the static clonal barcode, one circos plot per mouse. Concentric rings give the cell count, the sample group (tissue of origin), and the clonal barcodes; chords link clones shared between the primary tumor and the metastatic sites, so the number of shared clones reports the clonality of each site. (d) Global proliferation rank of clones (lambda or λ; rank 1 = fastest-growing). Point size scales with the clone’s frequency in that condition. In vivo frequencies are computed from the per-cell clonal barcodes. (e) Frequency-weighted mean lambda for each condition, against the mean of the fastest-growing 10% of clones (dashed line).

**Supplementary Figure 3.**
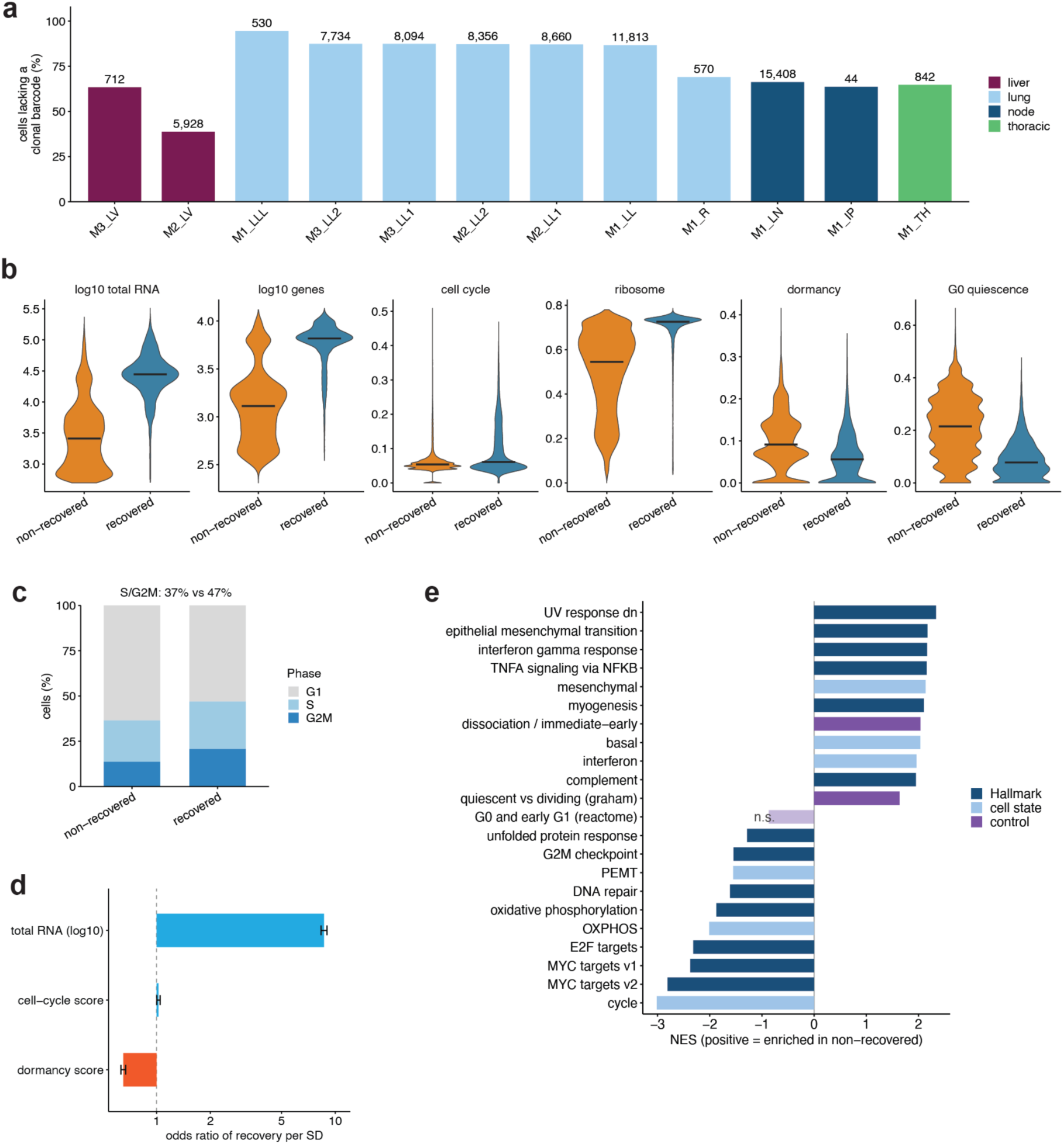
A pervasive low-recovery population is transcriptionally low-output and non-proliferative. (a) Fraction of transcriptomically defined tumor cells lacking a recovered clonal barcode, per sample (mouse and tissue; cell number above each bar), colored by tissue class. Non-recovery is lowest in the Mouse 2 liver metastasis (39%) and highest in the lung (87% to 95%), with the lymph nodes, thoracic mass, right lung, and the much smaller Mouse 3 liver metastasis intermediate (63% to 69%). The liver value is therefore driven by Mouse 2; the 712-cell Mouse 3 liver sits with the intermediate group. (b) Recovered (blue) versus non-recovered (orange) cells for total RNA and detected genes (log10) and the cell-cycle, ribosome, dormancy, and G0-quiescence UCell scores. Non-recovered cells are low-output and low-ribosome. Per-cell rank-based scores are depressed by dropout in low-complexity cells, so the pseudobulk enrichment in (e) is the primary readout. (c) Cell-cycle phase composition: non-recovered cells are less often in S or G2M (37% versus 47%). (d) Logistic model of recovery (odds ratio per standard deviation, tissue-adjusted): recovery is dominated by transcriptional output (total RNA, odd ratio 8.6 per SD), while a higher dormancy score predicts lower recovery once output is adjusted for (odd ratio 0.65) and the cell-cycle score is flat (odd ratio 1.02). (e) Module enrichment of the low-recovery population (primary characterization; same method as Figure 5b,c): fgsea on the depth-matched paired pseudobulk ranking against the Hallmark (dark) and pan-cancer cell-state (light) collections, with a dissociation/immediate-early set and two published quiescence references scored as pathways in the same run for direct comparison (purple; n.s. marks a control that is not enriched); NES > 0 = enriched in non-recovered. Proliferation (cell-cycle module, *MYC* and *E2F* targets, G2M), DNA repair and OXPHOS fall; Hallmark EMT, the mesenchymal module, interferon and inflammatory programs rise, while the narrower pEMT module falls. Of the two quiescence controls, the quiescent-versus-dividing reference rises (NES 1.64), but the mechanistic G0 and early-G1 pathway is not enriched (NES –0.87, adjusted P = 0.72), so the cell population is non-proliferative without showing a G0 entry program. The dissociation control also rises, which is why the claim rests on the falling axis. Abbreviations: NES, normalized enrichment score; UMI, unique molecular identifier.

For each cell, two barcodes quantify the event: the static clonal barcode reports clonality, and the number of independent clones seeding a site from the clonal sharing between primary and metastatic organs (**Figure 2c**); the evolving TAPE reports phylogeny, and from the shared edits among related cells, how many migrations founded a metastasis and roughly when (**Methods**; quantified per mouse in **Figures 3** and **Figure 4**). Together they place the three mice across a continuous range, from a single migration to the liver in Mouse 2 and Mouse 3 to hundreds of migrations converging on the lymph nodes in Mouse 1. Because both measurements require a recovered barcode, we restricted all clonal and phylogenetic analyses to barcode-recovered cells; the 77% of transcriptomically defined tumor cells that yielded none are a transcriptionally low, non-proliferative cell population rather than technical failures in barcode capture (**Supplementary Figure 2**), so the lineage reconstructions below exclude these cell population of each tumor (**Supplementary Figure 3**; **Supplementary Note 1**).

**Figure 3.**
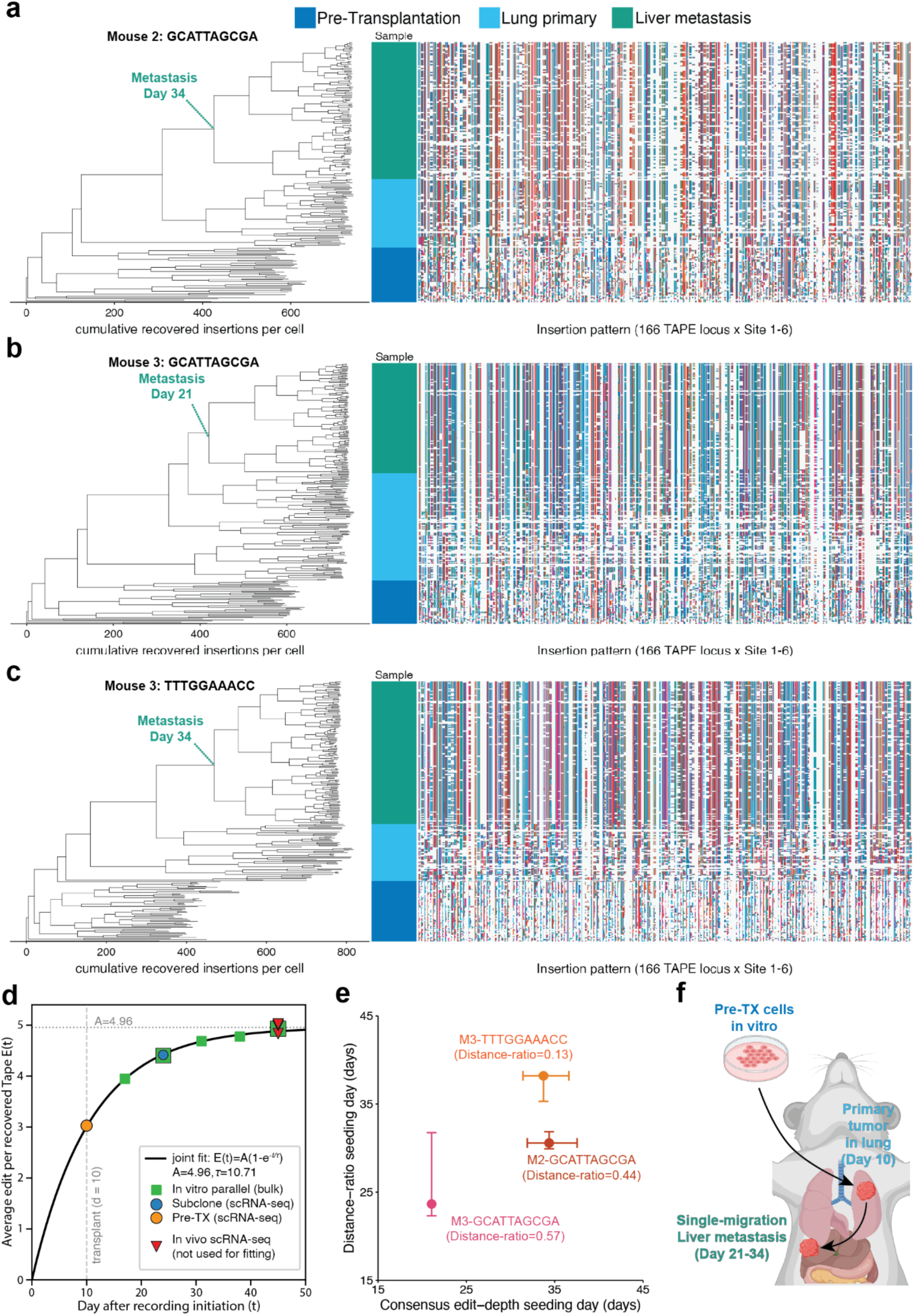
A calibrated single-cell recorder clock dates the dissemination of individual liver metastases in Mouse 2 and Mouse 3. (a-c) Edit-scaled single-cell lineage trees for the three liver-metastatic clones: (a) GCATTAGCGA-M2 (153 tumor cells and 41 pre-transplant), (b) GCATTAGCGA-M3 (208 tumor cells and 41 pre-transplant), and (c) TTTGGAAACC-M3 (229 tips, 176 tumor cells and 53 pre-transplant). (d) Recorder-clock calibration using the cumulative TAPE editing profiled at different time/modality. The single-exponential function, E(t) = A(1 – e^(–t/tau)), is fitted into single-cell time points and a rescaled bulk-amplicon series (fit: A = 4.96, tau = 10.71). (e) The liver-dissemination day from the consensus edit depth is compared with an estimate from the pairwise distance ratio (R; **Methods**). Vertical error bars are the distance-ratio cell-bootstrap 95% confidence interval; horizontal error bars are the clock-parameter (A, tau) bootstrap 95% confidence interval on the consensus-depth dissemination day (**Supplementary Figure 4b-d**). (f) Schematic of the single-migration liver-metastasis model in Mouse 2 and Mouse 3: pre-transplant cells expanded in vitro are transplanted and colonize the primary lung forming a tumor, from which a single founder cell disseminates to seed the liver metastasis (11 to 24 after the transplantation).

**Supplementary Figure 4.**
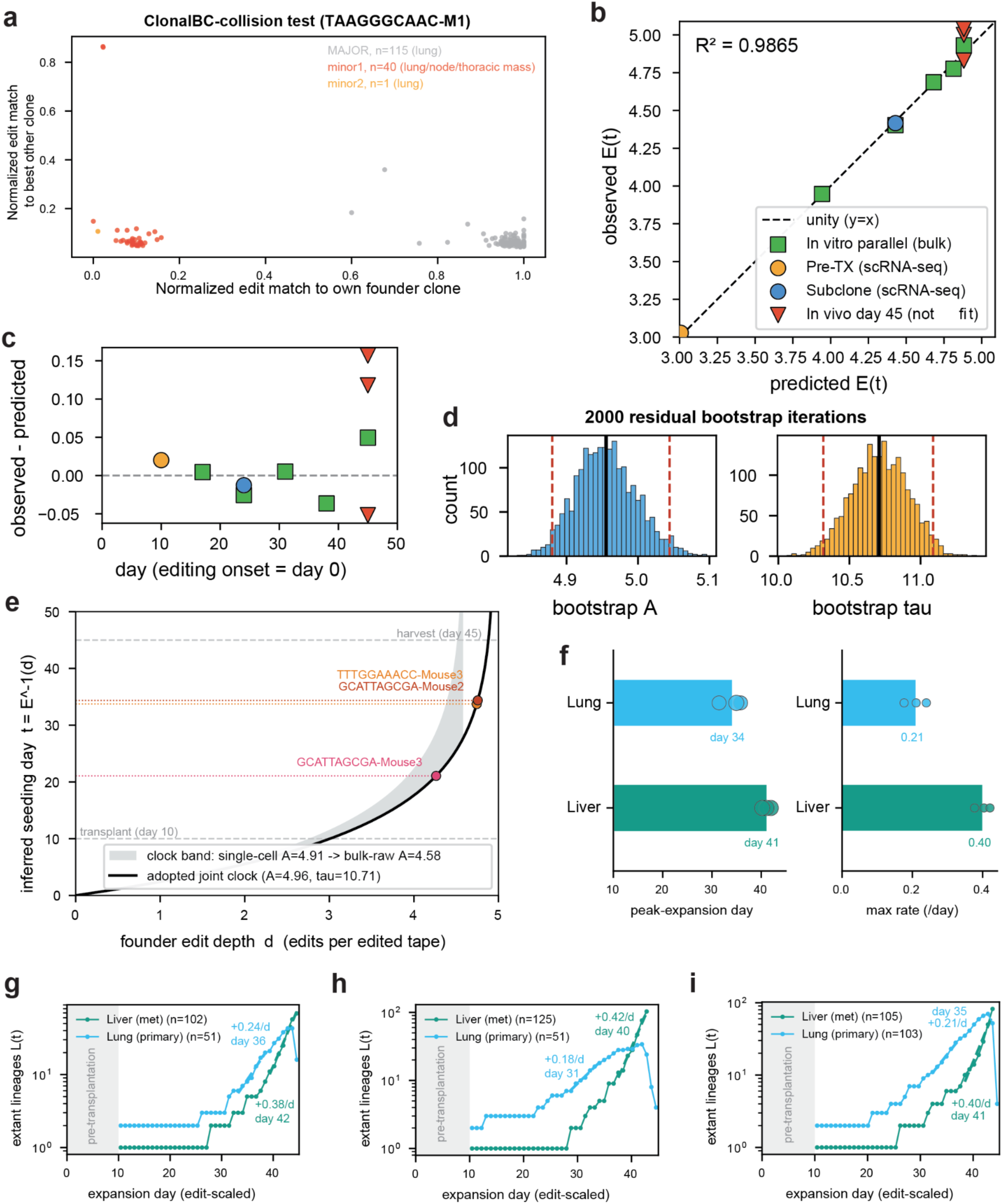
Founder screening, lineage growth, and recorder-clock calibration. (a) Founder and barcode-collision screen, which distinguishes a genuinely polyclonal barcode from a monoclonal seeding. The polyclonal example TAAGGGCAAC (Mouse 1) separates into a lung-restricted family (grey, 115 cells) and a distinct lymph-node and thoracic mass family (red, 40 cells) that share only the static clonal barcode, plus a single further cell assigned to neither (orange). (b) Clock goodness-of-fit: observed versus predicted edit depth E(t) across the bulk in-vitro calibration series, the two single-cell anchors shown separately (Pre-TX at day 10 and Subclone at day 24), and the held-out in-vivo day-45 cells (R^2 = 0.99). Colours match the dataset series of Figure 3d. (c) Calibration residuals (observed minus predicted E(t)) versus day, with the same series and colours as (g). (d) Bootstrap distribution of the fitted clock parameters over 2,000 residual bootstrap iterations, with tau refitted holding the bulk series fixed. Solid black rule, the point estimate; dashed red rules, the 95% confidence interval. A = 4.9553 (95% CI 4.8799 to 5.0443) and tau = 10.7089 (95% CI 10.3167 to 11.0886), so the calibration is well-constrained. (e) Dissemination-time sensitivity to the clock, confirming that the two near-saturation clones are robustly late. (f) Per-tissue lineage growth. Left, the day of peak lineage expansion for each tissue on the edit-scaled tree, whose depth is anchored linearly from transplantation (day 10) to harvest (day 45); the lung primary precedes the liver metastasis in every clone, with no overlap (lung mean day 34.1, range 31 to 36; liver mean day 41.1, range 40 to 42). Right, the maximum instantaneous expansion rate per tissue (lung mean 0.21 per day, liver 0.40 per day). Both quantities are invariant to a constant sampling fraction, so the comparison is robust to the primary being under-sampled relative to the liver. The order, not the absolute rate, is reported, because the axis orders expansion rather than dating it and the near-saturation liver clade is compressed on the edit-scaled axis. (g-i) Per-clone lineage-through-time curves underlying panel (f), one per dated clone. GCATTAGCGA-M2 in (g), GCATTAGCGA-M3 in (h), and TTTGGAAACC-M3 in (i).

**Figure 4.**
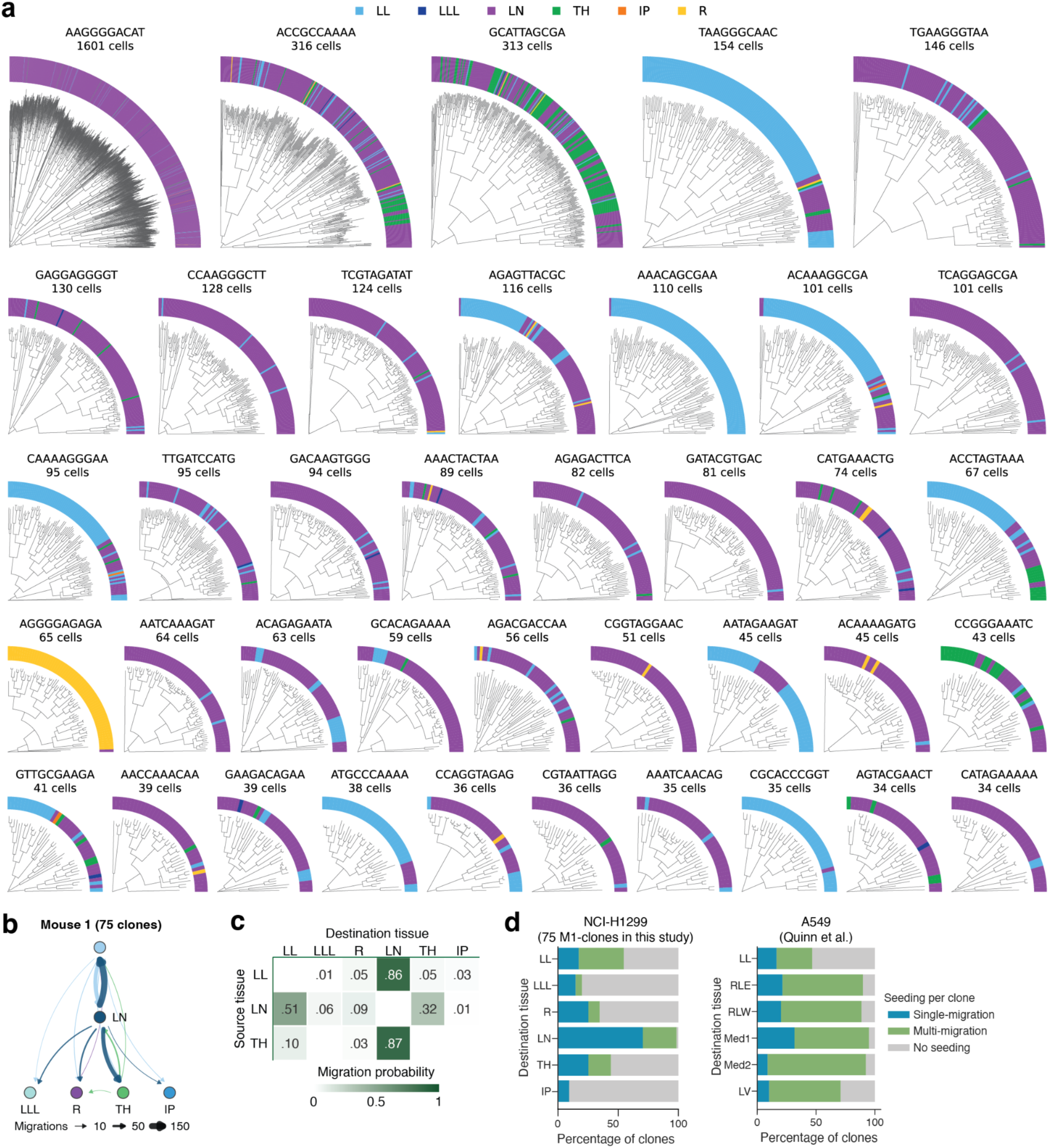
A lymph-node hub built from many migrations organizes polyclonal dissemination. (a) The 39 out of 75 Mouse 1 clonal lineages recovered with at least 34 cells, each a radial edit-scaled single-cell lineage tree (branch length in insertions gained) ordered by cell count. The outer ring colors each tip by sample location (LL, primary lung; LLL, lower-left lung; LN, mediastinal and regional lymph node; TH, thoracic mass; IP, intraperitoneal lymph node; R, right lung). (b) Aggregated inferred-migration graph across the 75 Mouse 1 clones (Metient): per-clone migration graphs are probability-weighted and summed. Nodes are anatomical sites and directed edges are inferred migrations, with edge width proportional to the number of migrations. The dominant route is from the primary lung (LL) to the lymph nodes (LN), which then redistribute cells to the lower lung (LLL), thoracic mass (TH), and intraperitoneal nodes (IP). (c) Tissue-transition matrix. Each entry is the probability that a migration leaving the source tissue (row) arrived at the destination tissue (column). Rows are normalized to migrations leaving each source and drawn from the most probable Metient solution per clone. LLL, R and IP are omitted as sources, as they emitted 0, 1 and 0 migrations respectively. (d) Per-clone seeding modality across destination tissues: (left) 75 NCI-H1299 clones from Mouse 1 in this study; (right) the A549 lung-adenocarcinoma xenografts from Quinn et al.^6^. The percentage of clones that seeded each tissue through a single migration (blue), through multiple migrations (green), or not at all (grey). LL, left lung; RLE/RLW, right lung; Med1/Med2, mediastinum 1/2; LV, liver.

We next asked whether proliferation drives colonization. We estimated each clone’s in vitro proliferation rate (lambda) from its abundance across the parallel in vitro time course (**Methods**) and placed the clones recovered in each tissue against a global ranking by lambda (**Figure 2d**). The clones that dominated the lung primaries and metastases were not the fastest growers but were spread across the full range of proliferation rank. Summarized as the frequency-weighted mean lambda per condition (**Figure 2e**), the in vitro pool drifted toward the fastest-growing decile over successive passages, as expected when faster clones outgrow slower ones, whereas every in vivo tissue across the three mice sat near the neutral pool mean and well below that band. Lung colonization and metastatic seeding are therefore not selected by intrinsic proliferation rate.

These three mice thus span the range we observed, from focal to widespread, consistent with lymphatic and distant metastases differing in the width of their evolutionary bottleneck^14^. We dissect the two ends in turn: first Mouse 2 and Mouse 3, whose focal liver metastases resolve into single founding cells and can be dated precisely, establishing the lineage-dating framework on unambiguous events, then Mouse 1 with a high-migration, lymph-node-centered dissemination.

### A recorder clock dates the liver metastases as late, single-founder events

Mouse 2 and Mouse 3 formed focal liver metastases with four liver-seeding clones recovered(**Figure 2b**). Three clones carried both primary-lung and liver cells and could be reconstructed as single-cell trees, while a fourth (ATGATCATAT-M2), recovered only in the liver, could not be dated against a primary. We reconstructed each tree from the ordered TAPE insertions under a sequential parsimony criterion that respects the irreversible order of editing, scaling branches by edits gained (**Methods**). Each metastasis is highly likely to be a single lung-to-liver migration founded by one cell, rather than a polyclonal or multi-migration event (**Figure 3a-c**, **Supplementary Figure 4a**).

To time these disseminations, we modeled the recorder’s editing dynamics from bulk and single-cell data, which followed the expected single-exponential accumulation^22^ across the 166 TAPEs, and used the fit to convert a consensus editing depth to an experimental date (**Methods**; **Figure 3d**; **Supplementary Figure 4b-d**). Dating each metastasis by its founder’s edit depth placed GCATTAGCGA-M3 earlier than others, at about day 21, and TTTGGAAACC-M3 and GCATTAGCGA-M2 near day 34 (**Figure 3a-c**). Because dissemination times near saturation are sensitive to the clock and the tree reconstruction (**Supplementary Figure 4e**), we cross-checked them with a second, largely model-free estimator from the ratio of liver-to-liver versus liver-to-primary cell distances, which is nearly independent of the editing rate (Distance-ratio; **Methods**). The two estimators agreed, to within half a day for the well-resolved early clone and on a late dissemination for the two near-saturation clones, and the distance ratio alone reproduced the ordering (*R* = 0.57, 0.44, 0.13; GCATTAGCGA-M3 earliest, TTTGGAAACC-M3 latest) without invoking the clock (**Figure 3e**).

Two further analyses confirmed that the liver lesions post-date the primary tumor. First, per-tissue lineage growth peaked later in the liver than in the lung in every clone (about day 41 versus day 34, with no overlap), so the metastasis both establishes and expands after the primary (**Supplementary Figure 4f-i**); the transcriptionally-low, non-proliferative cell population was correspondingly smaller in the Mouse 2 liver metastasis than in any lung sample (39% versus 86 to 95%; **Supplementary Figure 3a**), though the smaller Mouse 3 lesion is intermediate. Second, day-10 pre-transplant (Pre-TX) cells sharing each clone’s barcode grafted entirely basal to the lung-liver divergence (**Figure 3a-c**), separated cleanly from cells recovered in vivo. Together these place the liver metastases as genuine post-transplant seedings founded by single cells, not pre-existing subclones carried in with the xenograft (**Figure 3**).

### A lymph-node migration hub drives dissemination

These rare, precisely dated disseminations in Mouse 2 and 3 define one end of the metastatic range, analogous with the narrow lineage bottleneck described for distant metastases in human tumors^14,15^. We now turn to the opposite extreme in Mouse 1, where cells disseminated widely across the thoracic cavity rather than through single founders.

To reconstruct the overall pattern of metastasis, we reconstructed each clone’s migration history with Metient, which infers the tissue of origin at every tree node and counts migrations between sites^23^, rooting each phylogeny at the primary so that incomplete sampling of the primary tumor did not inflate migration counts (**Methods**). Among the 75 clones recovered with at least ten cells, individual trees routinely interleaved primary left-lung (LL) cells with lymph-node and other thoracic cells (39 out of 75 trees with at least 34 cells are shown in **Figure 4a**), showing that clones left the primary independently of one another rather than through a single dominant lineage.

Aggregated across clones, the migrations formed a strongly asymmetric, lymph-node-centred pattern (**Figure 4b,c**). Dissemination out of the primary was almost entirely directed to the mediastinal and regional lymph nodes: Of the 33 clones with an inferred migration out of the primary, 32 seeded the mediastinal and regional lymph nodes. The lymph nodes were in turn the most common origin of onward movement: 51 of 75 clones disseminated from LN, against 33 from the primary and 6 from any other site, and accounted for 76% of all inferred migrations. From the lymph nodes, clones reached the primary again (39 clones), the thoracic mass (28 clones), the right lung (20 clones), the lower left lung (14 clones) and the intraperitoneal nodes (4 clones). More than half of all clones (41 of 75) showed at least one migration back into the primary, almost all of them from LN. Nearly every clone reached the lymph nodes (99%), predominantly through a single migration (71%), whereas the distal sites were reached by far fewer (9-44%; **Figure 4d**). The lymph nodes thus acted as a central hub rather than a stop on a linear cascade, assembled from many independent single-migration seedings converging from across the population. This wide LN bottleneck against the narrower distal one parallels the distinction in human tumors between polyclonal lymph-node metastases and the narrower bottleneck of distant metastases^14^.

This lymph-node hub recapitulated the polyclonal, multidirectional seeding reported in a Cas9-based lineage-tracing study of lung adenocarcinoma xenografts^6^, in which every surveyed site was likewise seeded by many clones. Compared with that A549 cohort, our NCI-H1299 model showed fewer migrations per clone, a smaller multi-migration fraction, and more clones that seeded nothing at all (**Figure 4d**); the shared topology argues that lymph-node-centered, multi-migration dissemination is a reproducible feature of aggressive lung adenocarcinoma xenografts, while the lower migration rate let the opposite, single-migration extreme be sampled in Mouse 2 and Mouse 3.

Against patient data, the convergent LN seeding agrees with the clinical picture, in which lymph-node involvement is an early, prognostically significant, and characteristically polyclonal event^13^. The reseeding arm is more distinctive as our reconstruction inferred substantial LN-to-distal migration, whereas phylogenetic studies of human carcinomas attribute most distant metastases to direct seeding from the primary^12,15,24^. We consider the interpretation of this difference in the Discussion.

Taken together with the single-founder liver seedings of Mouse 2 and Mouse 3, these reconstructions span dissemination from one migration to more than a hundred, out of a single transplanted pool. Metastatic spread in this model is therefore not a fixed property of the population the cells come from, and any cell-intrinsic contribution to it should be expected to shift a clone’s odds of metastatic spread.

### Metastatic cells converge on a reproducible liver-adaptation program

Having established the origin of the liver metastases, we asked how the progeny of the founder cells adapt to their new environment. For each of the three clean liver metastatic clones (GCATTAGCGA-M3, TTTGGAAACC-M3, and GCATTAGCGA-M2), we assembled a clonally matched triad of single-cell transcriptomes from the pre-transplant in vitro pool, the lung primary, and the liver metastasis, so that transcriptional change is read within a clone and not confounded by clone-specific cell state. With a fourth clone added that was recovered in the liver but not the lung (ATGATCATAT-M2), cells separated jointly by environment and clone, the dominant axis distinguishing the in vitro pool from the two in vivo sites (**Figure 5a**).

**Figure 5.**
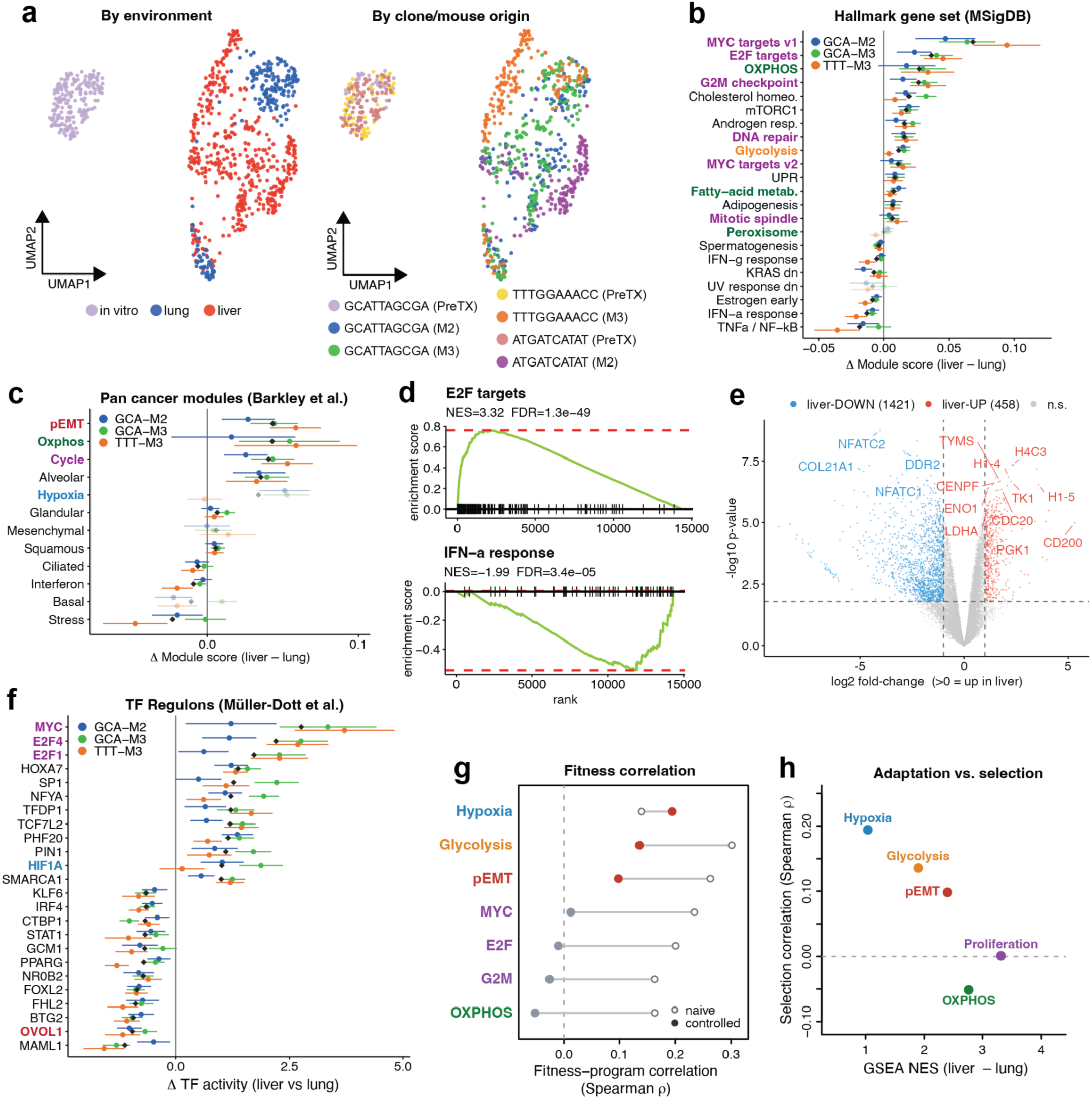
Metastatic cells converge on a liver-adaptation program that is decoupled from selection. (a) UMAP of the four clones across the three environments, colored by environment (left), colored by clone and mouse of origin (right). (b) Clonally resolved Hallmark gene sets^26^ with significant enrichment (FDR<0.05) in liver adaptation. For each gene set, the 95% confidence intervals from the per-cell scores are calculated for three clones, with a gray diamond indicating the across-clone mean. Programs whose liver-lung shift has the same sign in all three clones are drawn at full opacity. (c) Clonally resolved Pan-cancer gene modules^25^ in liver adaptation, as in **Figure 5b**. (d) Representative enrichment plots for the liver adaptation: *E2F* (top) and interferon-alpha response (bottom). (e) Volcano plot for the lung-to-liver using pseudobulk differential expression (5% FDR and at least two-fold change). (f) Clonally resolved regulator activity in liver adaptation, as in **Figure 5b**. (g) Correlation of transcriptional programs with phylogenetic fitness with (filled circles) or without (open circles) controlling for library size, mitochondrial content, and cell-cycle score. (h) Scoring each program by their enrichment in liver adaptation and selection, where a dashed line marks zero selection.

**Supplementary Figure 5.**
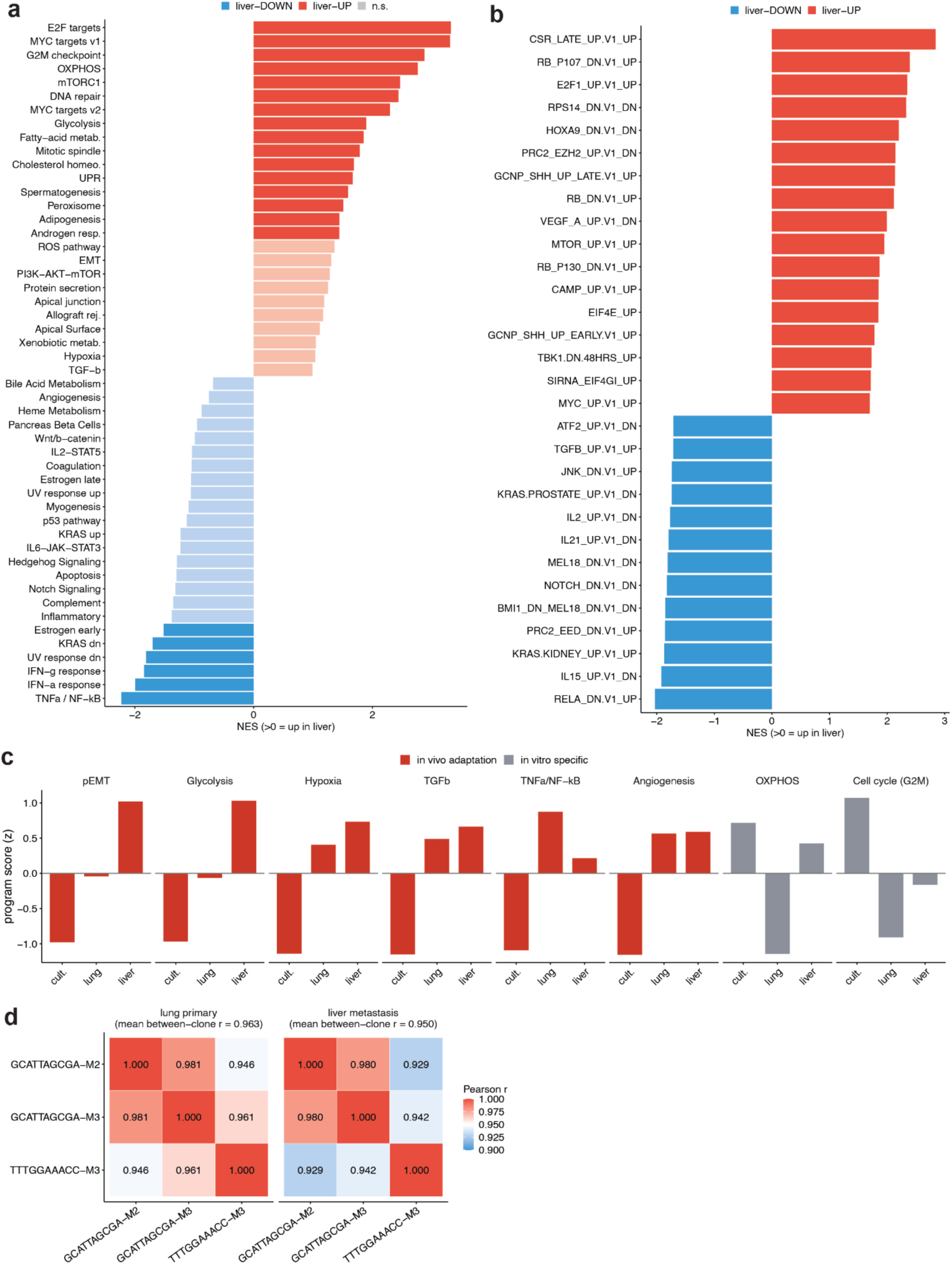
Extended liver-versus-lung enrichment, the three-environment trajectory, and between-clone similarity. (a) All 50 Hallmark gene sets ranked by normalized enrichment score (NES) for the liver-versus-lung contrast, from the pooled paired pseudobulk ranking (fgsea; NES > 0 = enriched in the liver). Bars are colored by direction and shaded by significance; 22 of the 50 sets reach a 5% false-discovery rate. (b) The 30 most extreme oncogenic (C6) signatures^26^ of the 189 tested, ranked by NES for the same contrast. The RB-loss and E2F-activation signatures corroborate the proliferative axis of **Figure 5b**. (c) Culture to lung to liver program trajectory, one small multiple per program. Bars are the mean program score (UCell^31^) averaged over the three genotype-matched clones and z-scored across the three environments, and are colored by trajectory shape: red, programs induced in vivo relative to culture, including the liver-adaptation programs; grey, programs highest in culture that fall in the lung and recover only partly in the liver. (d) Between-clone transcriptome similarity by environment (Pearson correlation over 2,000 highly variable genes), for the lung primary (left) and the liver metastasis (right). Clones remain transcriptionally distinct in both tissues, and the liver metastases are no more similar to one another than the primaries are (mean between-clone r = 0.963 in the lung, 0.950 in the liver).

**Supplementary Figure 6.**
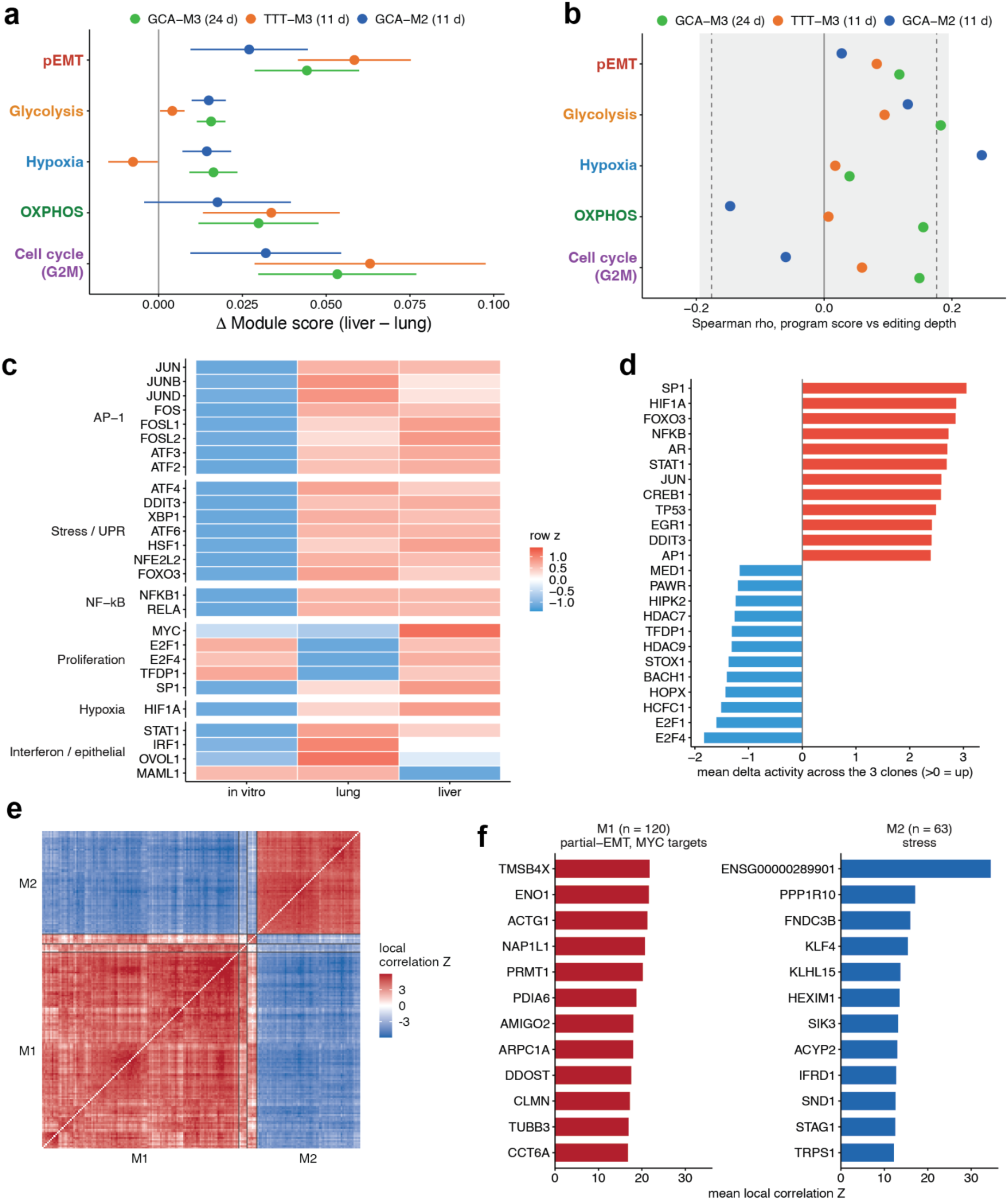
Residence time and the transcription-factor regulators of the liver-adaptation program. (a) Liver-minus-lung difference in each of the five liver-adaptation programs, one series per clone, with 95% confidence intervals from the per-cell scores. Clones are labelled with their residence time in the liver at harvest (harvest day minus the dissemination day of **Figure 3**). The longest-resident metastasis, at 24 days, is not the most adapted in any program, and the two seeded about 11 days before harvest reach the same or larger differences, so the program is installed rapidly rather than deepening over this window. (b) Internal control for (a). Within each liver metastasis, the per-cell Spearman correlation between each program’s score and that cell’s TAPE-recorder editing depth (332 liver cells over three clones). The shaded band is the two-sided 5% significance range, 1.96/√(n − 1), drawn as a range because n differs between clones; the dashed lines mark its narrowest extent. (c) Curated regulon activity across the three environments (row z-score), grouped by family (AP-1, stress and unfolded-protein response, NF-kB, proliferation, hypoxia, interferon and epithelial). Cell means are averaged within each clone first and then across clones, to avoid the domination of the liver column by the single 3,387-cell clone (ATGATCATAT-M2). (d) Data-driven regulon activity of the culture-to-lung entry step (decoupleR^27^ over the CollecTRI network^28^), showing the twelve transcription factors rising and the twelve falling most by across-clone mean activity. (e) Cross-clone consensus of the lineage-informed co-expression structure in the three liver metastases. Hotspot was run per clone with that clone’s edit-scaled single-cell lineage tree as the cell-to-cell metric; each clone’s local correlation matrix was then averaged across clones and clustered, over the 194 genes recurrently autocorrelated among them. Blocks on the diagonal are the four consensus modules, of which the two large ones are labelled (M1, 120 genes; M2, 63). Module numbering is independent of the in-vitro consensus of **Figure 6a**. Colour is the averaged local correlation Z, clipped at its 99th percentile; the diagonal is blank because self-correlations are excluded. (f) The twelve strongest genes of each of the two large consensus modules of (e), ranked by mean local correlation Z, with the programs that module is enriched for shown above it (one-sided hypergeometric over the 194 genes entering the consensus). The two anti-correlated blocks of (e) are an adaptation and biosynthesis module set against a stress module (7 of 7 Barkley stress genes, *P* = 3 × 10^-4^). The adaptation module is enriched for partial-EMT (14 of 14 set members present, *P* = 9 × 10^-4^) and *MYC* targets (19 of 22, *P* = 0.008), and carries the cytoskeletal and glycolytic genes *TMSB4X*, *ACTG1* and *ENO1*. The strongest gene of the stress module, *ENSG00000289901*, is a 218-base unnamed lncRNA on chromosome 13 with no assigned symbol; it is recovered independently in all three metastases but is not in the heritable set of **Figure 6** and is null in the pre-transplant outcome-prediction screen in **Figure 6**.

We scored each cell from the three metastatic clones for pan-cancer cell-state modules^25^ and MSigDB Hallmark gene sets^26^ and tested liver-versus-lung shifts by pseudobulk differential-state analysis with the clone as replicate (**Methods**). Because the pre-transplant pool was captured separately, we will focus our quantitative analysis only on the lung-to-liver comparison (a shared capture and harvest), and interpret the culture-to-lung change at the qualitative changes in program directions.

Comparing each liver metastasis with its paired lung primary resolved a reproducible program in which proliferative and metabolic axes led. Among Hallmark modules, *MYC* and *E2F* targets were the two strongest signals (Normalized Enrichment Score or NES = 3.3 each), followed by oxidative phosphorylation (OXPHOS; NES = 2.8), mTORC1, and glycolysis (NES = 1.9), with interferon and TNF-α-NF-κB programs coordinately down (**Figure 5b**, **Supplementary Figure 5a**). The pan-cancer curated modules^25^ added a partial epithelial-to-mesenchymal transition (pEMT; NES = 2.4, FDR = 7 × 10⁻¹¹) and a hypoxic program, both up in the liver, with the cell-cycle module the most strongly enriched of that collection (NES = 3.7) (**Figure 5c**). The two reciprocal shifts are shown as running-enrichment plots (**Figure 5d**): the rising proliferative programs (*E2F*, with *MYC* and G2M alongside; all FDR < 1 × 10⁻²⁹), corroborated by oncogenic *RB*-loss and *E2F*-activation signatures (**Supplementary Figure 5b**), against the falling interferon-α response (NES = −1.9). Together these enrichments nominate five programs for the liver adaptation: proliferation (cell cycling), OXPHOS, glycolysis, pEMT, and hypoxia.

At the gene level, pseudobulk differential expression identified 4,669 genes at 5% FDR, of which 1,879 changed at least two-fold (**Figure 5e**, **Supplementary Table S3**). The liver-up genes were led by replication-dependent histones and S-phase and mitotic genes (*H4C3*, *TK1*, *CENPF*) and a coordinated glycolytic block (*ENO1*, *PGK1*, *LDHA*); NFAT-family and stromal-matrix genes (*NFATC1/2*, *DDR2*) led the liver-down genes. Because these interferon and TNF-α-NF-κB programs are growth-suppressive and the host is immunodeficient, their decrease most likely reflects the reciprocal of the proliferative rise rather than an immune interaction.

This program is the far end of a directional culture-to-lung-to-liver trajectory rather than an isolated endpoint: the same liver-up programs, with Notch and the unfolded-protein response, rose monotonically from culture to lung to liver in all three clones (each still significant at the lung-to-liver step, FDR < 1 × 10⁻⁵), whereas a separate general in-vivo response (TGF-β, TNF-α-NF-κB, angiogenesis) distinguished culture from host irrespective of organ (**Supplementary Figure 5c**). Proliferation was the exception, highest in the rapidly dividing culture, lowest in the lung, and only partly reactivated in the liver, a liver-specific reactivation rather than a monotonic feature of metastasis.

Because all cells were harvested on the same day, the three dated disseminations span a residence-time axis in the liver (about 24 days for GCATTAGCGA-M3 versus 11 days for the two late clones). The adaptation did not deepen over this window: all three metastases were similarly adapted relative to their paired lung, the most recently arrived, if anything, the most adapted, so the program is installed rapidly and essentially complete within about ten days of dissemination (n = 3; **Supplementary Figure 6a,b**). The convergence is program-specific, where the liver metastases were no more globally similar to one another than the primaries, and clones remained transcriptionally distinguishable in the liver (**Supplementary Figure 5d**). Therefore, cells converge on the shared modules while keeping clonal identity.

### Regulator and lineage analyses independently recover the liver-adaptation program

To identify the regulators, we inferred transcription-factor activity with decoupleR^27^ over the curated transcriptional factor regulons^28^. Entry into the host raised a broad immediate-early and stress program (AP-1, integrated-stress and unfolded-protein regulators, and *NF-κB*, among others; **Supplementary Figure 6c**), a step that is batch-confounded as noted above (**Supplementary Figure 6d**); we therefore focused on the within-capture lung-to-liver change, dominated by rising *MYC*, *E2F*, and *HIF1A* activity and falling *OVOL1* (epithelial), *MAML1* (Notch), and *STAT1* (interferon) activity, recovering the proliferative, pEMT, and hypoxic axes (**Figure 5f**).

To find programs organized by lineage rather than transcriptional similarity alone, we ran Hotspot^29^ with the single-cell lineage tree as the cell-to-cell metric. Averaged across the three metastases, the recurrent lineage-informed modules resolve into two anti-correlated consensus modules (**Supplementary Figure 6e**): a partial-EMT and *MYC*-target module that carries the cytoskeletal and glycolytic genes (*TMSB4X*, *ACTG1*, *ENO1*), and a stress module (**Supplementary Figure 6f**). Glycolysis and hypoxia appear here as individual genes within the first module rather than as enriched sets; because a metastasis seeded by a single cell is itself a monophyletic clade, this whole-tumor signal coincides with the culture-to-liver transition and cannot on its own separate inherited from environmentally induced expression, so the direct heritability test is deferred to the next section. Together, these pathway, gene-level, regulator, and lineage-informed analyses converge on the same five programs as the reproducible signature of liver adaptation (**Supplementary Table S4**).

### Fitness is decoupled from the strongest liver adaptation

We next asked which of these programs are under selection in the liver metastases themselves, using the single-cell lineage trees of the metastatic clones. Estimating a phylogenetic fitness for each cell from the shape of its lineage tree (the local branching index^30^), higher-fitness lineages naively scored higher on nearly every program: proliferation, OXPHOS, partial-EMT and hypoxia alike. Controlling for library size, mitochondrial content and cell cycle collapsed the proliferative (*MYC*, *E2F*, G2M) and OXPHOS associations to zero or below in every clone, while hypoxia strengthened (0.14 to 0.19) and glycolysis and partial-EMT weakened but stayed positive (0.30 to 0.14 and 0.26 to 0.10; **Figure 5g, Supplementary Table S5**). The fitness-correlated lineages are therefore not simply the most proliferative, but carry a hypoxia-, glycolysis– and partial-EMT-centered state. Because tree resolution limits power and the local branching index partly reflects graft expansion, the ordering among programs is more robust than the absolute correlations (**Methods**).

Plotting each program by its liver-adaptation magnitude against its association with fitness separates what a metastatic cell most strongly adapts from what is actually selected (**Figure 5h, Supplementary Table S6**). The two are inversely related across all five programs: proliferation and OXPHOS are the most strongly upregulated in the liver yet carry no selection signal, whereas the selected programs are the more modestly upregulated hypoxia, glycolysis and partial-EMT. What a metastatic cell upregulates most strongly on arrival is therefore largely a plastic response to the liver, whereas the smaller hypoxia-, glycolysis– and partial-EMT-centered axis correlates with selection.

### Transcriptional basis of the heritable metastatic potential

The transcriptional basis of the heritable metastatic potential must satisfy two requirements: First, the expression of involved genes must be inherited during the culturing prior to transplantation. Second, clones with higher expressions of these genes at the time of transplantation must predict their initial colonization of the primary injection site as well as their metastasis. To reveal the candidate genes for the heritable metastatic potential, we tested both requirements directly and broadly, then asked which ones satisfy them together.

To test heritability of each gene in culture, we used eight monoclonal colonies expanded in vitro from the pre-transplant pool, each grown from a single founder and profiled so that a high-resolution lineage tree and the transcriptome came from the same cells (295 to 11,081 cells per colony). Using Hotspot^29^ on each clonal tree, we calculated lineage autocorrelations as heritability and grouped them into gene modules, whose module scores were sharply structured along the lineage (**Figure 6a**, left, a representative colony; **Supplementary Figure 7**). Across the eight different colonies, 396 genes showed high and recurrent heritability, arranged into five broadly annotated modules (**Figure 6a**, right). While Hotspot uses a fixed neighbourhood size to calculate heritability, phylogenetic autocorrelation can be also calculated using PATH^32^ (Phylogenetic Analysis of Transcriptional Heritability; **Methods**) that uses all pairs of the tree tips. The two agree closely (Spearman rho = 0.86; **Supplementary Figure 8a**), and we took their consensus as the pre-existing heritable set of 349 genes (**Figure 6b**, **Supplementary Figure 8b**, **Supplementary Table S7**). We then asked whether the previous five liver-adaptation programs were present in this heritable gene set: partial-EMT was the most strongly enriched program (36 of 96, 10.8-fold), followed by proliferation (36 of 186, 5.6-fold), hypoxia (4.6-fold) and glycolysis (2.8-fold), with OXPHOS barely above chance (1.6-fold) (**Figure 6c**).

**Figure 6.**
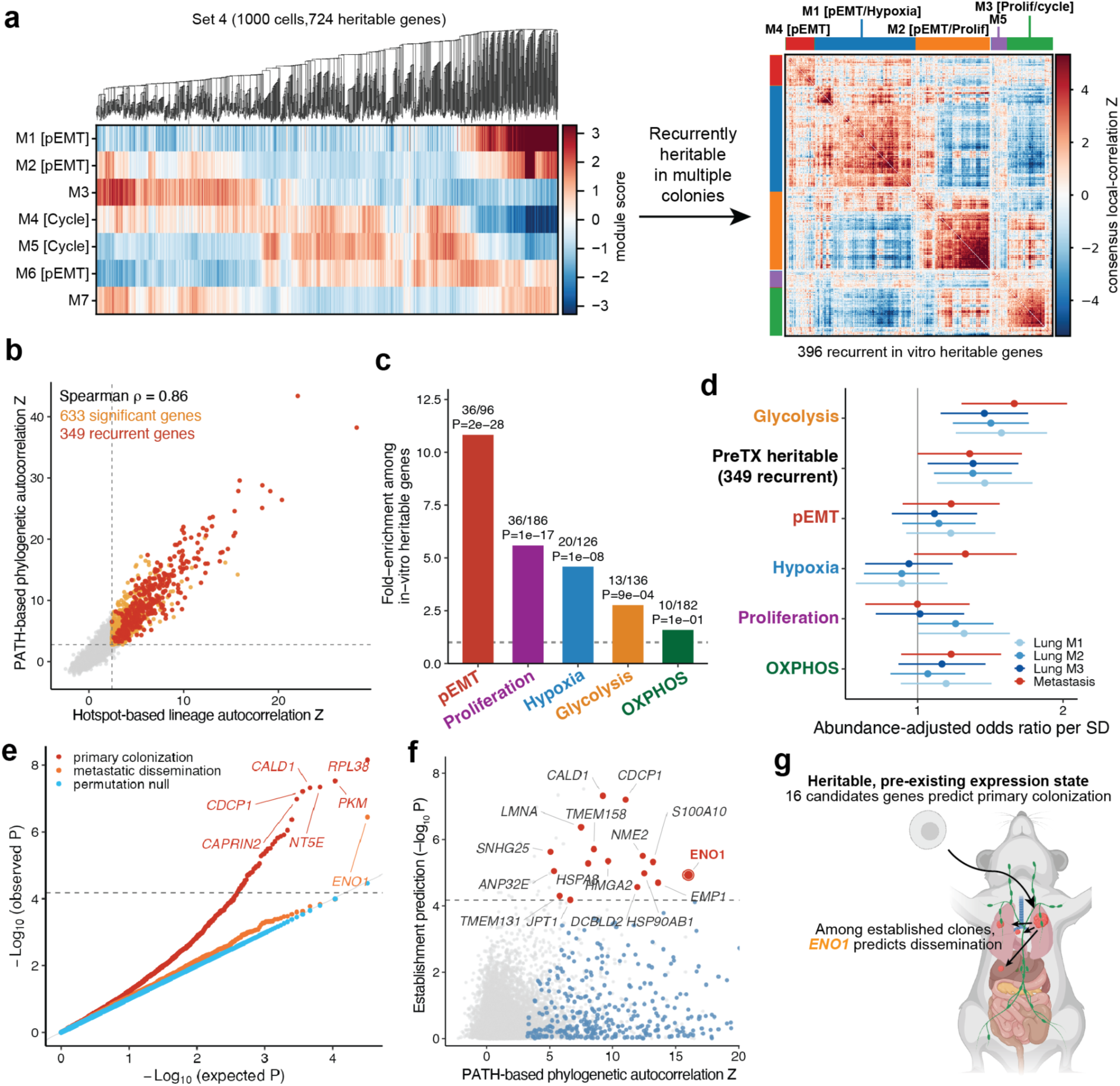
A pre-existing glycolytic state predicts which clones colonize and disseminate. (a) Calculating heritability of each gene in vitro. Left: a lineage tree of representative monoclonal pre-transplant colony (Set 4), and module scores for 7 detected gene modules plotted below. Right: the consensus gene-by-gene lineage co-expression matrix over all 396 genes called heritable by Hotspot in at least 75% of the colonies. (b) Heritability measurements using Hotspot versus PATH. The 633 genes above both threshold but failing the recurrence criterion are colored in orange, while the 349 gene recurrently heritable in at least 75% of the colonies by both methods are colored in red. (c) Program enrichment in the heritable gene set (one-sided hypergeometric, over the 349 genes). (d) Per-program logistic models over the 1,229 pre-transplant clones with at least 10 cells, each adjusted for the clone’s abundance in the pool. Points are the odds ratio per standard deviation (95% CI); series are colonization of the primary lung in each of Mouse 1, 2 and 3 (blue) and dissemination (red), the latter comparing clones recovered at a secondary site against clones that established but remained in a lung. (e) Quantile-quantile plot of association P values over all 16,221 detected genes against two clone-level outcomes, establishment (red) and dissemination among established clones (orange). The background from 200 rounds of reshuffling the outcome labels across clones colored in blue. (f) The 10,065 genes categorized by their lineage heritability (mean PATH Z) versus evidence for establishment (*−log₁₀ P*). The 349 heritable genes are colored blue, the 16 of those that also predict establishment at 5% FDR (dashed line) are colored red, and *ENO1* is highlighted with a ring. (g) Schematic of the model where a heritable, pre-existing expression state resolves to sixteen genes that individually predict establishment, and among established clones that have already been established, *ENO1* alone predicts dissemination.

**Supplementary Figure 7.**
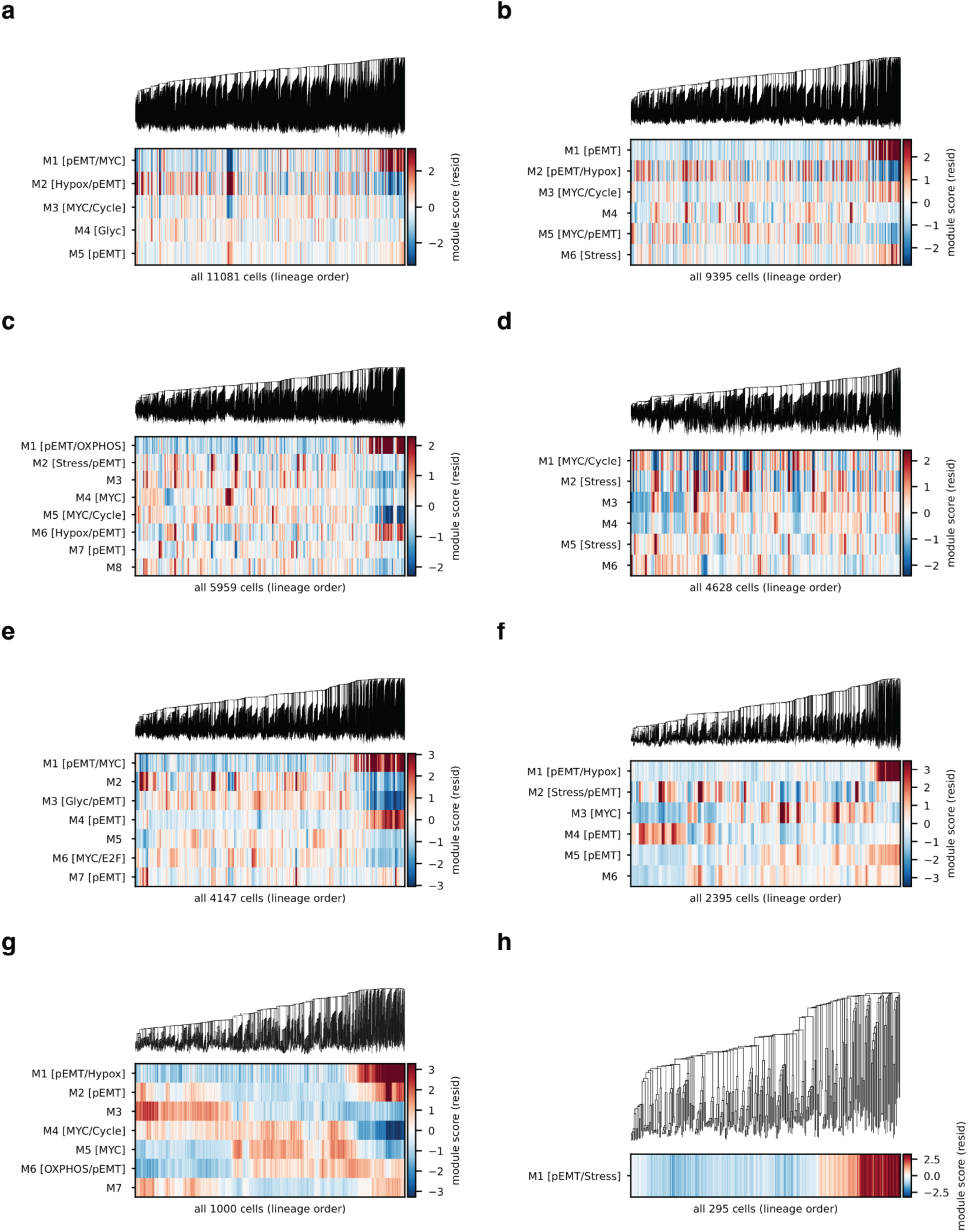
Lineage heritability in each of the eight monoclonal pre-transplant colonies. (a–h) One panel per colony, ordered by cell number: the full-depth Cassiopeia lineage tree above a heatmap of per-cell heritable-module scores in lineage order, residualized on mitochondrial fraction and log total counts so that a block along the tree is not a depth or quality gradient. Colonies contain 295 to 11,081 cells and yield 86 to 2,283 lineage-autocorrelated genes (Hotspot FDR < 0.05). Module numbering and annotation are assigned independently within each colony and do not correspond between panels, nor to the consensus modules of **Figure 6a**. The smallest colony (h, 295 cells) is power-limited and resolves a single module, which is partial-EMT annotated. Panel (g) is the colony shown in **Figure 6a**.

**Supplementary Figure 8.**
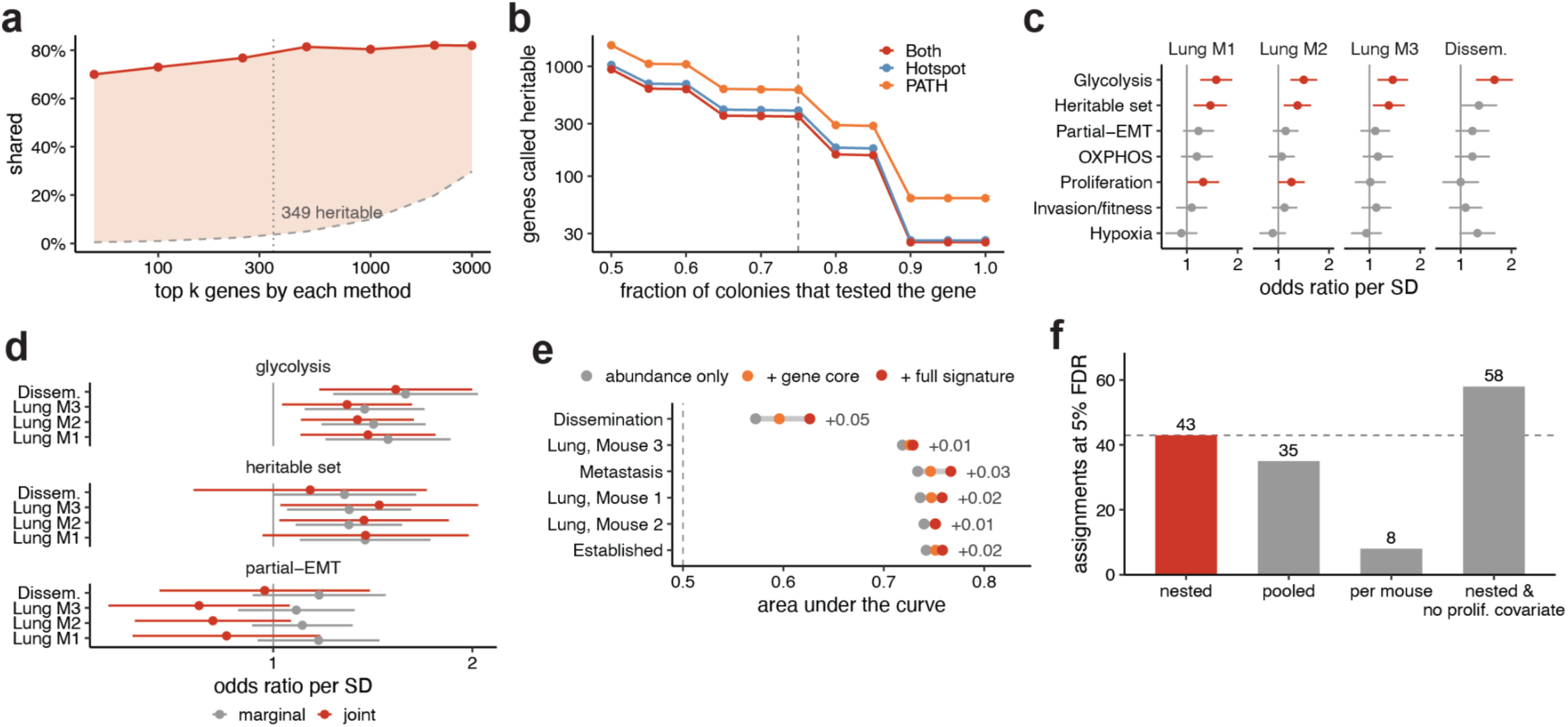
Robustness of the heritability, prediction and gene-scan results of Figure 6. (a) Overlap of the top k genes ranked by mean Hotspot Z and by mean PATH Z, over the 10,078 genes scored by both. Red, observed; dashed grey, chance overlap at that k; shading, the excess over chance. Dotted line, the 349-gene heritable set. (b) Number of genes called heritable against the recurrence fraction, for each method and their intersection, requiring at least four colonies tested throughout. Dashed line, the 75% used in the main figures. (c) All signatures fitted alone and adjusted for pool abundance, across the three primary-lung outcomes and dissemination. Red, *P* < 0.05. Proliferation reaches nominal significance in Mouse 1 and Mouse 2 but not Mouse 3 and not for dissemination, with both confidence intervals bounded at one. (d) Odds ratio per standard deviation for the three competence signatures, each adjusted for pool abundance. Grey, each fitted alone; red, all three fitted together. Glycolysis and the outcome-independent heritable set both retain their effect on colonization under joint adjustment. Partial-EMT inverts below one in every contrast. (e) Area under the curve for each outcome from models containing pool abundance alone (grey), abundance plus a compact gene core (14 heritable glycolysis gene of *CITED2, ENO1, FUT8, HDLBP, LDHA, MIF, P4HA1, PAM, PGK1, PKM, PPIA, SOX9, TPI1, TXN*; orange), and abundance plus the full signature (191 MSigDB glycolysis set^26^; red); the segment spans abundance alone to the full model and the number is the gain. Each mouse’s primary lung is tested separately. “Established” is recovery anywhere versus never. “Metastasis” includes any secondary sites excluding left lung, and “Dissemination” compares metastasis with lung-only clones, excluding clones never recovered. (f) Gene-outcome assignments passing 5% FDR under four outcome-labelling schemes; red, the nested scheme used in the main figure (dissemination nested in the established clones), with the dashed line at its count of 43.

Next, we asked whether the pre-existing heritable gene set predicts which clones colonize the primary lung and which then disseminate. Throughout, colonization refers to recovery in a given mouse’s primary lung and is scored per mouse; establishment refers to recovery anywhere in any mouse against recovery nowhere, pooled across mice. Scoring each of the 1,229 pre-transplant clones with at least 10 cells (31,838 cells) for every program and for the set itself, we tested first which clones were later recovered in the lung, controlling for abundance in the pool, and then, among those that established, which were also found at a metastatic site. Among six programs tested, a pre-existing glycolytic state and heritable gene set predicted colonization of the primary lung reproducibly in all three mice (*P* < 0.003 and *P* < 0.02, respectively) (**Figure 6d**, **Supplementary Figure 8c**). Hypoxia was null here despite carrying the strongest fitness association within the metastases (**Figure 5g**), as expected for a state that is selected in vivo but induced by the niche rather than present beforehand. Conditioned on establishment, glycolysis was the only one of the signatures tested that also predicted dissemination (1.59, *P* = 3 × 10^-4^), while the heritable set was borderline (1.28, *P* = 0.051). Fitted together, glycolysis and the heritable set both retain their effect on colonization while partial-EMT inverts below one and reaches significance nowhere, so the two are separable rather than one restated, and both shift a clone’s odds modestly against a recovery process dominated by its abundance in the pool (**Supplementary Figure 8d,e, Supplementary Table S8**).

Finally, we tested each of the 16,221 detected genes individually for association with establishment and with metastatic dissemination against a background generated by reshuffling the outcome labels across clones. Forty-three genes were above a 5% false-discovery rate for establishment, and *ENO1*, a glycolytic enzyme that also moonlights as a cell-surface plasminogen receptor^9^, was the only one also above a 5% false-discovery rate for metastatic dissemination (**Figure 6e**; **Methods**). Dropping the proliferation covariate does not reduce the number of significant assignments (**Supplementary Figure 8f**). Comparing these with the recurrently heritable gene set, sixteen candidates emerged (**Figure 6f**): *CALD1, CDCP1, LMNA, TMEM158, SNHG25, NME2, HMGA2, S100A10, HSPA8, ANP32E, HSP90AB1, EMP1, DCBLD2, TMEM131, JPT1* and *ENO1*. The two requirements are near-independent across genes (Spearman rho = 0.12). Two of them, *ENO1* and *S100A10*, are also cell-surface plasminogen receptors.

Taken together, the basis of heritable metastatic potential in this model is a transcriptional state already organized along the lineage before transplantation, not a difference in clonal growth rate (**Figure 2e**). It acts in two steps: a glycolytic set-point and an outcome-independent lineage-heritable set separately predict whether a clone colonizes the primary lung, while among the clones that establish only the glycolytic axis predicts onward dissemination (**Figure 6g**). At gene level, sixteen genes are both recurrently heritable in culture and predictive of establishment, of which *ENO1* alone also predicts following dissemination. Because the scan detects only effects large enough to clear a genome-wide threshold across 1,229 clones, these sixteen are a lower bound on the contributing genes rather than a complete account.

## DISCUSSION

To date, several lineage recorders have been used to study cancer metastasis^6,7,16,33^, but resolving each metastatic event has been difficult because the recordings are unordered insertions and deletions that saturate the recording sites quickly. By engineering an ordered, high-capacity DNA Typewriter lineage recorder in NCI-H1299 cells and measuring its editing rate at multiple time points across the 45-day experiment, we dated disseminations that seeded individual liver metastasis to specific days (day 21 to 34, i.e., 11 to 24 days after transplantation). The same recorder, read across three mice, differed in their dissemination pattern depending on their host. Focal liver metastases were each found by a single cell, while a high-rate of dissemination can be observed through a lymph-node as a transitional hub. Yet, neither pattern was selected by intrinsic proliferation rate observed in vitro.

Profiled across three mice, the reconstructed lineage trees revealed two dissemination patterns that recapitulate observations from endpoint human cancer genomics. In Mouse 2 and Mouse 3, focal liver metastases were each founded by a single cell arriving directly from the lung primary, matching both the narrow, late bottleneck reported for distant metastases from lung and the finding that most distant metastases seed directly from the primary rather than from lymph node deposits^12,15,24^. Mouse 1 instead disseminated at a high rate within the thoracic cavity and adjacent nodes, through a lymph-node hub with substantial onward migration to the other thoracic sites, consistent with the wider evolutionary bottleneck reported for lymph node metastases^13,14^ and with reseeding through nodal vessels^34^. However, we cannot exclude that a portion of the earliest regional spread in Mouse 1 was seeded at or shortly after orthotopic injection rather than by later migration, a caveat specific to this model and to sites near the injection. The clones dominating both patterns were spread across the full range of in vitro growth rates, so neither was driven by intrinsic proliferation rate.

Joint analysis of the reconstructed lineage trees with the transcriptome let us ask what state predisposed cells to seed and how they adapted in the new tissue environment. Metastatic cells converged on a reproducible liver-adaptation program across genotype-matched clones, led in magnitude by proliferation and OXPHOS. However, the programs upregulated most strongly on arrival were not those under selection. Controlling for cell quality and cell cycle, phylogenetic fitness within the metastases correlated not with proliferation or OXPHOS but with a smaller hypoxia-, glycolysis– and partial-EMT-centered axis, and of the programs tested only a pre-existing glycolytic state and the lineage-heritable set predicted which clones later colonized the primary lung in vivo. Metastatic competence in this model is therefore a pre-existing, heritable cell state rather than a proliferative advantage, present in the founding cells before they encounter the host. This gives a molecular basis for the classical observation that metastatic variants pre-exist within a tumor without a simple mutational cause^5^, and it agrees with recent evidence that carcinoma cells leave the primary tumor with predetermined metastatic potential^35^.

The heritable metastatic potential resolves into two separable components: a glycolytic program and the broader set of lineage-heritable genes, each predicting which clones colonize the primary lung in the host. We treat this potential as a probabilistic risk factor that shifts a clone’s odds rather than a deterministic switch, given the differences in metastatic pattern and frequency across mice. In our screen, only *ENO1* predicted which of the established clones went on to disseminate, and whether that reflects its glycolytic role or one of its moonlighting functions is a natural question. In lung cancer, *ENO1* overexpression raises glycolysis, migration and invasion in vitro and metastasis in vivo, with knockdown reversing both^36^. More notably, *ENO1* also has a non-glycolytic function as a plasminogen receptor^9^, where antibody blockade of the surface pool of *ENO1* alone suppresses matrix degradation and metastasis^37^. The strongest adhesion candidate in our scan, *S100A10*, also has a role as a plasminogen receptor^38^. Notably, the previous lineage-tracing study of pancreatic cancer found the *S100* family associated with metastatic subpopulations^7^. A shared plasminogen-receptor function would link the glycolytic and adhesion components that our models otherwise treat as separable, a possibility worth testing both for its role in metastasis and as a therapeutic entry point, as urokinase plasminogen activator receptor (uPAR) has recently been targeted by CAR T cells in preclinical models^39^.

Whichever function *ENO1* acts through, the glycolytic component remains the dominant heritable predictor of primary lung colonization, consistent with the large literature tying aerobic glycolysis, the Warburg effect, to tumor progression and metastasis^40,41^. Metabolic requirements along the cascade are stage-dependent, however: reports that metastasis depends on OXPHOS, from PGC-1α-driven mitochondrial biogenesis in invading breast-cancer cells^42^ to an OXPHOS-dominant phenotype in circulating tumor cells^43^, concern cells already in transit or re-initiating growth, whereas we identify a pre-dissemination state: a heritable glycolytic state present in normoxic culture and in the founder cells, and therefore cell-intrinsic rather than hypoxia-driven, that marks which clones later establish. A model of glycolytic priming before dissemination followed by metabolic flexibility during transit and outgrowth accommodates both. It remains an association, however, and separating a causal dependency from a correlated marker will require direct measurements and glycolytic perturbation of defined clones, which this lineage-resolved system is well placed to enable.

Several limitations bound these conclusions: Because the recorder is read out by amplicon capture that scales with transcriptional output, it cannot reach the large transcriptionally-low, non-proliferative cell population that pervades every site (**Supplementary Figure 3**; **Supplementary Note 1**), so the lineage reconstruction here is limited to the actively proliferating cells. Whether those cells are truly dormant rather than simply non-cycling is a second axis of heterogeneity for future work with an improved system that captures genomic recordings from cells with low RNA counts. Next, the timed disseminations rest on three reconstructable liver clones (n = 3), and more recovery of metastatic clones with recent migration is needed to resolve adaptation to the liver environment fully. Finally, the model is a single lung cancer line (NCI-H1299) in immunodeficient hosts, so tumor-immune interaction is absent.

More broadly, this work shows how a structured lineage-recording experiment can recover the biology that endpoint sampling misses. Transplanting a single pool with clonal barcodes and molecular recorders into several hosts and recovering genomic recordings and transcriptome from the same cells reveals when a metastasis seeded, from which founder, and in what state, separating a cell’s pre-existing competence from what it acquires on arrival. Here that heritable metastatic potential reduces to a compact, directly assayable state: a heritable glycolytic component that predicts which clones colonize and, among those that establish, which disseminate. Whether the same cell state governs metastatic potential in immunocompetent and genetically engineered models and in human tumors will be the natural next test. The dissociated readout used here loses tissue context, so pairing the recorder and transcriptome with spatial or in-situ methods^16,44^ would be more powerful still, placing a founder’s state and its lineage within the niche it colonizes. The notion of heritable metastatic potential is complementary to the high-plasticity cell state that drives progression and therapy resistance in autochthonous lung tumors^45,46^, in that the state they describe is defined by its capacity for transition whereas the competence we identify is stably transmitted along the lineage. This separation of heritable from plastic contributions, and the molecular recording methods that resolve it, are well-poised to extend beyond metastasis to other settings of tumor adaptation such as acquisition of new cancer traits and drug resistance^47,48^.

## ENDNOTE

## ACKNOWLEDGEMENTS

We thank the members of the Choi and Kim labs for helpful discussions, and Jay Shendure and Tuomas Tammela for feedback and advice throughout the project. This research was supported, in part, by NIH/NCI Cancer Center Support Grant P30CA008748, by the Basic Science Research Program through the National Research Foundation (NRF) of Korea grants funded by the Korean government (MSIT) (RS-2022-NR070713 and 2018R1A5A2025079 to H.H.K.), the NRF of Korea grants funded by the Ministry of Education (RS-2023-00248758 to J.P.), the Bio and Medical Technology Development Program of the NRF funded by the Korean government (MSIT) (RS-2022-NR067326 and RS-2023-00260968 to H.H.K.), the Korea Drug Development Fund funded by the Ministry of Science and ICT, the Ministry of Trade, Industry and Energy, the Korea-US Collaborative Research Fund (KUCRF), the Ministry of Science and ICT and Ministry of Health & Welfare, Republic of Korea (RS-2024-00467177 to H.H.K.), the Brain Korea 21 FOUR Project for Medical Science (Yonsei University College of Medicine), the Yonsei Fellow Program funded by Lee Youn Jae, and the Samsung Research Funding & Incubation Center of Samsung Electronics (SRFC-MA1701-51 to H.H.K.). D.K. is supported by the National Science Foundation Graduate Research Fellowship (227260-01) and NIH (5T32GM132083-02). J.Chen is supported by the NIH (K08CA259161) and American Lung Association (LCD-1252549). D.A.L. is supported by the NIH Common Fund Somatic Mosaicism Across Human Tissues UG3NS132139, National Human Genome Research Institute, Center of Excellence in Genomic Science RM1HG011014, Burroughs Wellcome Fund Career Award for Medical Scientist, Vallee Scholar Award, Blood Cancer United Scholar Award, and the Mark Foundation Emerging Leader Award. This work utilized resources from the High Performance Computing Group at Memorial Sloan Kettering Cancer Center. J.Choi is supported by the NIH (R00HG012973), Damon Runyon Cancer Research Foundation (DFS-64-24), and Searle Scholars Award (SSP-2025-101).

## AUTHOR CONTRIBUTIONS

J.P., Y.C., H.H.K., and J.Choi initiated the project.

J.P. and Y.C. performed experiments with inputs from J.S., H.H.K., and J.Choi.

J.P. and J.Choi analyzed the data with inputs from J.S., H.S., C.M., J.Chan, Q.M., and D.A.L.

J.S. developed the Distance-ratio method.

D.K. performed Metient analysis with inputs from J.P., Q.M., and J.Choi.

J.Choi and H.H.K. supervised the study.

J.P. and J.Choi wrote the manuscript with input from all authors.

## METHODS

### Plasmid design and cloning

The lineage-recording system comprised three integrated components. The prime editor (PE) construct encoded a Cre-inducible PE2max prime editor (“PB-EF1a-loxP-HygR-3xPolyA-loxP-PE2max-P2A-mClover3”): a loxP-flanked HygR-pA(x3) cassette positioned upstream of PE2max such that Cre-mediated recombination excises the cassette and activates PE2max expression, with mClover3 as a fluorescent reporter. The pegRNA-Tape library construct encoded a pegRNA together with an mRFP-marked recorder array of six Tape monomers (6x Tape) flanked by a Read2N primer-binding site and a CS1 sequence and carried a 10 nt static TargetBC (N10) identifying each recorder Tape cassette. The ClonalBC library construct encoded a Puromycin-resistance marker (PuroR) and luciferase, together with a 10 nt static clonal barcode (ClonalBC, N10) flanked by Read2N and CS1 sequences.

Both the ClonalBC and pegRNA-Tape constructs were cloned by Gibson assembly (NEBuilder HiFi DNA Assembly Master Mix, E2621, NEB). Inserts were generated by an overlap-extension approach: each was ordered as a pair of forward and reverse oligonucleotides (Macrogen) whose 3’ ends share ∼10–25 bp of complementarity and which additionally carry homology arms to the vector for Gibson assembly. For each insert, the two oligonucleotides (50 pmol each) served simultaneously as primers and templates: their complementary 3’ ends anneal and are mutually extended by Q5 High-Fidelity DNA polymerase (M0491, NEB) to generate the full-length double-stranded insert, under the following conditions: 98 C for 30 s; 15 cycles of 98 C for 30 s, 60 C for 20 s, and 72 C for 30 s; and a final extension at 72 C for 5 min.

*ClonalBC plasmid library.* The amplicon generated with primers “ClonalBC_N10_fwd” and “ClonalBC_N10_rev” was used as the insert. The backbone vector (“pLenti-EF1a-Puro-P2A-Luciferase-WPRE”) was digested with MluI-HF (R3198, NEB) at 37 C for 2 h, treated with Quick CIP (M0525, NEB) at 37 C for 30 min, and gel purified (28506, Qiagen) before the Gibson assembly reaction. After incubation at 50 C for 30 min, the Gibson assembly mixture was purified by isopropanol precipitation and transformed into TransforMax EC100 Electrocompetent E. coli (EC10010, Biosearch Technologies). Bacterial colonies were harvested, and the final plasmid library (“pLenti-EF1a-Puro-P2A-Luciferase-ClonalBC-WPRE”) was extracted using a Plasmid Maxi Kit (12165, Qiagen).

*pegRNA-Tape plasmid library.* The pegRNA-Tape plasmid library was generated by a two-step cloning process. First, to clone the pegRNA region for “NNNNGGA” insertion, the insert with “Tape_peg_NNNNGGA_fwd” and “Tape_peg_NNNNGGA_rev” was generated as described above. The backbone vector (“PB-U6-pegRNA-BbsI-BbsI-EF1a-mRFP-TAPE-BsmBI-BsmBI”) was digested with BbsI-HF (R3539, NEB) at 37 C for 2 h and treated with Quick CIP at 37 C for 30 min; the subsequent steps followed the ClonalBC library protocol. This intermediate plasmid library was harvested and extracted using a Plasmid Maxi Kit. Second, the amplicon generated with “Tape_targetBC_N10_fwd” and “Tape_targetBC_N10_rev” was used as the insert to clone the TargetBC region, and the intermediate plasmid library was digested with BsmBI-v2 (R0739, NEB) at 55 C for 3 h and treated with Quick CIP at 37 C for 30 min; the subsequent steps again followed the ClonalBC library protocol. The final pegRNA-Tape plasmid library (“PB-U6-pegRNA-NNNNGGA-EF1a-mRFP-TAPE-TargetBC”) was extracted using a Plasmid Maxi Kit. The primers used for cloning are described in **Supplementary Table 1**.

### Lentivirus production

HEK293T cells (CRL-1573, ATCC) were cultured in Dulbecco’s Modified Eagle Medium (11995-065, Thermo Fisher Scientific) supplemented with 10% Fetal Bovine Serum (FBS, RDTech Inc.) and 1% pen-strep (15140-122, Thermo Fisher Scientific). To produce lentivirus, the Luciferase-ClonalBC transfer plasmid (“Lenti-EF1a-Puro-P2A-Luciferase-ClonalBC-WPRE”) was mixed with the packaging plasmids psPAX2 (Addgene; #12260) and pMD2.G (Addgene; #12259) at a 4:3:1 weight ratio, giving 60 µg of total plasmid. This mixture was transfected into HEK293T cells at 70–80% confluence in a 15 cm dish with Lipofectamine 2000 (11668-019, Thermo Fisher Scientific). The medium as replaced with 20 mL of fresh growth medium 24 h after transfection, and virus-containing supernatant was harvested 72 h after transfection and passed through a 0.45 µm Millex™ PVDF syringe filter (SLHV033NS, Millipore). Aliquots were stored at –80 C until use. Viral titer was determined as previously described^22^.

### Cell line and lineage recorder engineering

NCI-H1299 (KCLB 91299, Korean Cell Line Bank), a human non-small cell lung carcinoma line with loss of p53 expression and an activating NRAS Q61K mutation, was maintained in RPMI-1640 (11875-093, Thermo Fisher Scientific) with 10% fetal bovine serum, 1% sodium pyruvate, 1% HEPES and 1% penicillin-streptomycin at 37 °C in 5% CO2.

The DNA Typewriter recording system (Choi et al., Nature, 2022) was installed by piggyBac transposition. Cells were transfected with 0.8 to 1.2 µg of a Cre-activatable prime editor, 6 to 9 µg of a pegRNA-TAPE library carrying a 10-nucleotide static TargetBC per array, and 0.2 µg hyPBase^49^, then selected with 100 µg/mL hygromycin B for two weeks and single-cell sorted on high mRFP and absent mClover3. Each TAPE array carries six sites written directionally in strict 5-prime to 3-prime order, so a site can be edited only after the site preceding it is filled. All construct, array, barcode and primer sequences are in **Supplementary Table S1**.

Five monoclonal candidates were screened by bulk amplicon sequencing of the TargetBC region for array copy number and, with and without Cre, for recording efficiency and Cre dependence (**Supplementary Figure 1a,b**). The selected line, Founder-39, was then transduced with a luciferase-ClonalBC-puromycin lentivirus at a multiplicity of infection of 0.3 with 10 µg/mL polybrene and selected with 1 µg/mL puromycin for 10 days (day – 11; **Supplementary Figure 1c**), marking each founding cell with a 10-nucleotide static barcode. The static ClonalBC reports which founding clone a cell belongs to; the evolving TAPE array resolves the genealogy within that clone.

Recording was initiated at day 0, the time origin for every date reported here, by transfecting 1 µg Cre recombinase plasmid with Lipofectamine 3000 into 800,000 cells per well across six wells. On day 3, the top 30% of mRFP-positive, mClover3-positive cells were sorted and bottlenecked to approximately 8,000 cells, then expanded approximately 20-fold so that several siblings of each founding clone were represented. The pool was split four ways: approximately 10,000 cells orthotopically injected into the left lung of each of three mice at day 10; 40% profiled by single-cell RNA sequencing at transplantation (pre-transplantation pool, Pre-TX); an aliquot single-cell sorted into wells and expanded as monoclonal colonies; and the remainder maintained as a parallel in vitro culture with weekly bulk amplicon profiling to day 45.

### Animal experiments and tissue collection

All experiments conducted with live animals were approved by the Institutional Animal Care and Use Committee of Yonsei University Health System. Three female 8-week-old NOG mice (Central Institute for Experimental Animals) were used in this study. All mice were maintained under specific-pathogen-free conditions.

*Orthotopic injection of lineage-recording cells into mouse lungs.* NCI-H1299 cell suspensions were prepared for orthotopic implantation as follows. Cells were harvested from culture by trypsinization, and the reaction was quenched with growth medium. Following a wash in cold PBS, cells were resuspended in cold Matrigel matrix (A1413202, Thermo Fisher Scientific) at a final concentration of 1,000 cells/µL. The cell–matrigel suspension was gently mixed, loaded into a 1 mL syringe, and kept on ice until the time of implantation. Orthotopic implantation was performed as previously described^6^. Mice were positioned in the right lateral decubitus position and anesthetized with 2.5% inhaled isoflurane. A 1-cm incision was made along the posterior medial line of the left thorax, and the fascia and adipose tissue were dissected and retracted to expose the lateral ribs, intercostal space, and left lung. Once respiratory movement of the left lung was confirmed, a 30-G needle was inserted approximately 3 mm into the lung parenchyma through the intercostal space, and 10 µL of the cell suspension (5,000 – 10,000 cells per mouse) was injected directly into the lung tissue. Mice were monitored for 1-2 hours following the procedure, with body weight and wound healing assessed weekly thereafter.

*Sample dissociation for single-cell experiment.* Mice were euthanized by CO₂ inhalation, followed by exsanguination via transection of the abdominal aorta after laparotomy extending from the abdominal to the thoracic cavity. Fluorescent tumor lesions were identified and dissected under a stereomicroscope equipped with a fluorescence adapter, based on RFP signal (Nightsea). The upper left lung (LL), corresponding to the lobe used for initial tumor cell injection, was designated as the primary tumor site. Lower left lung (LLL), right lung lobes (R), mediastinal and regional lymph nodes (LN), thoracic mass (TH), intraperitoneal lymph node (IP), and liver (LV) were collected as metastatic sites when RFP-positive lesions were present, though the specific sites involved varied between individual mice. Single-cell suspensions were prepared as previously described^6^, with the following modifications. Unless otherwise noted, all steps were performed on ice. Tissue dissociation buffer was freshly prepared prior to each experiment using DMEM/F-12 medium supplemented with collagenase (3,000 U; LS005273, Worthington biochemical), Liberase TL (2.5 mg; 5401020001, roche), and DNase I (50 U; LS006344, Worthington biochemical) per 15 mL. Minced tumorous tissues were placed in gentleMACS C Tubes (130-093-237, Miltenyi Biotec) with 5 mL of dissociation buffer per tube and homogenized using a gentleMACS Octo Dissociator for 1 min. Tissues were then incubated in a 37 °C water bath, with manual trituration by pipetting performed every 5 mins (three to six cycles total) until fully dissociated. Cells were pelleted using a swinging-bucket rotor at 800 × g for 5 mins at 4 °C for all subsequent centrifugation steps. To lyse red blood cells, pellets were resuspended in ACK Lysing Buffer (1 mL per sample; A1049201, Thermo Fisher Scientific) and incubated for 3 mins at room temperature, followed by centrifugation. For single-cell dissociation, pellets were resuspended in 500 µL TrypLE reagent (12605010, Thermo Fisher Scientific) and incubated at 37°C for 5 mins; dissociation was assessed by pipetting, and incubation was extended in 5-min increments as needed for incompletely dissociated samples. Digestion was quenched with 1 mL of complete RPMI 1640 medium. Cells were centrifuged and resuspended in cold complete RPMI 1640 (final volume 300 – 500 µL depending on pellet size), then passed through 40 µm strainers. RFP-positive cancer cells were isolated by FACS sorting at 4 °C to exclude host-derived cells and debris; viability and sorting efficiency were monitored by flow cytometry. Sorted cells were pelleted (800 × g, 5 mins, 4 °C) and resuspended in PBS containing 0.04 % BSA to a final volume of 100 µL. For samples requiring further debris removal, an additional wash step was performed by resuspending cells in 1 mL of 0.04 % BSA/PBS, filtering through a 40 µm FlowMi filter tip (13680-0040, SP Bel-Art), and re-pelleting (800 × g, 5 mins, 4 °C) prior to final resuspension in 100 µL. Cell concentration was determined by manual counting.

### Genomic DNA extraction, and amplification of Clonal barcode and Tape sequences

Two sample types were processed: clonal screen samples and in vitro parallel samples. For clonal screen samples, 30 µL of direct lysis buffer containing 10 mM Tris-HCl pH 8.0 (15568-025, Thermo Fisher Scientific), 0.05% SDS (15553-027, Thermo Fisher Scientific) and 25 µg/mL proteinase K (EO0492, Thermo Fisher Scientific) was added to each well of a 96-well plate. Plates were incubated at 37 C for 1 h, followed by heat inactivation at 80 C for 30 min. Using 2 µL of the direct lysate as template, the Tape region was amplified with primers “Tape_amp_fwd” and “Tape_amp_rev” and PrimeSTAR GXL DNA Polymerase (R050B, Takara) under the following conditions: 95 C for 3 min; 33 cycles of 98 C for 10 s, 60 C for 15 s, 68 C for 3 min; and a final extension at 68 C for 5 min. The first PCR products were then indexed by 12 cycles of PCR and sequenced on the Illumina MiSeq platform.

For in vitro parallel samples, genomic DNA was extracted from each sample using the Wizard Genomic DNA Purification Kit (A1620, Promega) according to the manufacturer’s instructions. Using 2 µg of genomic DNA as input, the Tape sequence was amplified with primers “Tape_amp_fwd” and “Tape_amp_rev” and PrimeSTAR GXL Polymerase under the following conditions: 95 °C for 3 min; 23 cycles of 98 °C for 10 s, 60 °C for 15 s, 68 C for 3 min; and a final extension at 68 °C for 5 min. For the ClonalBC amplicon, 2 µg of genomic DNA was amplified with Q5 High-Fidelity DNA Polymerase (M0491, NEB) using primers “ClonalBC_amp_fwd” and “ClonalBC_amp_rev” under the following conditions: 98 °C for 30 s; 33 cycles of 98 °C for 10 s, 61 °C for 20 s, 72 °C for 1 min; and a final extension at 72 °C for 5 min. All amplicons were then indexed by 12 cycles of PCR and sequenced on the Illumina MiSeq platform. The primers used for PCR are described in **Supplementary Table 1**.

### Single-cell library preparation and sequencing

Sorted mRFP-positive cells were processed with GEM-X Single Cell 3’ Reagent Kits v4 (CG000731 Rev B, 10x Genomics). Feature cDNA Primer 3 replaced the standard cDNA primers so that the Read 2N primer was included and TAPE cDNA co-amplified, and the v3.1 purification (CG000389 Rev B, step 2.3) was used to separate three libraries by amplicon size: whole transcriptome (cell state), a TAPE recorder amplicon carrying its TargetBC (lineage), and a ClonalBC amplicon from the luciferase transcript (clonality). The ClonalBC library followed CRISPR Screening Library Construction and the TAPE library Cell Multiplexing Library Construction with PrimeSTAR GXL polymerase in place of the Amp Mix; the three were indexed with Dual Index TT, NT and NN kits respectively.

Gene expression and ClonalBC libraries were sequenced on Illumina NovaSeq 6000 and X Plus (R1/i7/i5/R2 = 28/10/10/90) and TAPE libraries on an AVITI24 (Element Biosciences; I1/I2/R1/R2 = 10/10/28/260), to median depths of approximately 20,000, 2,000 and 15,000 reads per cell respectively.

Sixteen libraries were generated: two pre-transplantation captures (Initial1, Initial2), two subclonal-colony captures (Subclone1, Subclone2), six Mouse 1 organ captures, and three each from Mouse 2 and Mouse 3. Across all samples, 105,260 single cells were profiled. Reads were aligned and demultiplexed with Cell Ranger v9.0.1 against human GRCh38, with the recording constructs added as custom reference features so that recorder-derived reads carry the feature tags ‘TAPE_TargetBC’ and ‘Luc_ClonalBC’ in the position-sorted BAM.

### Single-cell sequencing data pre-processing

*Extraction of clonal barcodes and Tapes from alignments.* Clonal barcodes and Tape edits were extracted directly from Cell Ranger BAM files using custom Python scripts (Step 1, Step 2 in analysis pipeline; see Data and code availability). Only reads carrying the appropriate fx feature tag and both a corrected cell barcode (CB) and corrected UMI (UB) were considered. For ClonalBC, the sequence between the two flanking sequences was recovered, and distinct UMIs counted per “cell barcode-ClonalBC” pair. For Tape, the cassette was isolated between its flanking sequences, the TargetBC was read from immediately after CATGCCCAAA, and the monomer array trimmed at TGATGGTGACTACATCTA. Monomer copies were located allowing up to 2 mismatches, and the sequences lying between consecutive monomers were recorded as the insertion pattern, distinct UMIs were counted per “cell barcode-TargetBC-insertion pattern-monomer count” combination. Barcodes deviating from the expected 10 nt length, and cassettes lacking an expected constant sequence, were flagged for downstream filtering. In both cases, only records supported by at least 2 distinct UMIs were retained.

*Clonal barcode assignment and error correction.* Raw ClonalBC counts were filtered and collapsed using a custom R script (Step 2c in analysis pipeline). ClonalBC sequences with a barcode length other than exactly 10 nt were discarded. Within each cell, ClonalBCs supported by fewer than 50% of the UMIs compared to that cell’s most abundant ClonalBC were removed. Each cell was then uniquely assigned the single ClonalBC with the highest UMI count. To correct for sequencing errors, clonal barcodes were collapsed based on a Hamming distance of 1. Distinct barcodes were ranked by their abundance (from most to least abundant). In a greedy clustering approach, each unassigned barcode was designated as a cluster centroid, and any less-abundant barcode within a Hamming distance of 1 was merged into it. Once a barcode was merged into a centroid, it could not act as a centroid for other barcodes; therefore, the merging process was restricted to a single level of depth and did not chain. The final, corrected ClonalBC list is provided in **Supplementary Table 2**.

*Doublet detection using Scrublet.* Doublets were identified per library with Scrublet^50^, run on the raw Cell Ranger count matrix with an expected doublet rate of 0.01, min_counts = 2, min_cells = 3, min_gene_variability_pctl = 85, and 30 principal components. Per-cell doublet scores and calls were exported for use during Seurat single-cell integration.

*Tape edit table construction.* Edit tables were assembled per group (Pre-TX (initial), Subclone, Mouse1, Mouse2, Mouse3) in Python. Within each group, per-sample Tape records were restricted to cells passing QC and doublet filtering, then pooled. Records were restricted to a curated set of 166 static TargetBCs (TargetBCs detected above a coverage threshold in the engineered clone screen and validated by amplicon sequencing). For each “cell barcode-TargetBC” pair, the insertion pattern with the highest UMI support was retained. Cells are kept for lineage reconstruction if they carry at least 100 retained TargetBCs (Initial and Subclone samples) or at least 20 TargetBCs (Mouse 1–3 samples. The resulting table was pivoted to a matrix with one row per cell and one column per TargetBC–site combination, with uninserted or undetected positions recorded as “None”.

*Single-cell quality control, normalization and integration.* Per-library matrices were processed using Seurat^51^ (v5.4.0) in R (v4.5.2). Genes detected in fewer than 3 cells were removed. Cells were retained if they expressed more than 200 and fewer than 15,000 genes and had less than 10% mitochondrial reads. Counts were log-normalized with a scale factor of 10,000. Cell barcodes were appended with their sample of origin, and libraries were merged into group-specific matrices (Initial (Pre-TX), Subclone, Mouse1, Mouse2 and Mouse3). Following normalization, cells flagged as doublets by Scrublet were removed from the merged objects. Cell-cycle phase was scored using the S and G2/M gene sets provided with Seurat, and the corresponding scores were regressed out during scaling. Highly variable features were identified, principal component analysis was performed, and the first 30 principal components were used for UMAP embedding and neighbor-graph construction. Clusters were called using the Leiden algorithm at a resolution of 0.3. The pre-implantation and mouse-derived samples were integrated into a single object. The subclone samples were processed separately through an identical pipeline and are presented as an independent embedding.

### Validation of lineage reconstruction using subclones

To assess whether the Tape lineage tree recovers true clonal relationships, we compared lineage tree clades against ground-truth clonal identities defined by a serial dilution assay in a 96-well plate. Ten wells of clonally expanded subclones were individually selected, dissociated, and pooled. The ClonalBC set carried by each well was established for 8 of 10 subclones; the remaining 2 clones had very low cell counts (14 and 22 cells) and were excluded from subsequent analysis.

*Lineage distance computation for reconstructed lineage validation.* To reconstruct lineage trees for the subclone quality-control analysis, pairwise distances between cells were computed from the Tape edit table. The distance metric was designed to reflect the ordered, cumulative nature of Tape editing, in which an insertion at a given site is only informative if all preceding sites in that cassette have also been edited. Within each Tape cassette, only a leading run of observed insertions was considered: sites following the first unedited (empty) position were treated as unobserved. For each pair of cells, similarity was defined as the total number of leading insertion sites that matched across all cassettes, counting sites in order and stopping at the first mismatch or unedited position within each cassette. The distance between cell *i* and *j* was then defined as

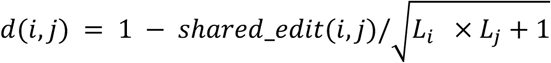

where *shared_edit(i,j)* is the number of matched edits summed across all sites between cell *i* and *j*, and *L_i_* is the total observed number of edits summed across all sites for cell *i*. This normalization scales shared editing by the geometric mean of the two cells’ recorded edit depths, so that cells with more extensive shared editing histories are placed closer together.

*Concordance between lineage tree and clonal barcode-based assignment.* Lineage relationships were reconstructed from the Tape edit distances. Using the pairwise distance matrix, cells were hierarchically clustered by average linkage (UPGMA), and the resulting tree was divided into 8 clades with *cutree*. In parallel, each cell was assigned to a well using its Hamming distance-corrected ClonalBC (top-UMI barcode per cell; see “Single-cell sequencing data pre-processing: Clonal barcode assignment and error correction”). A cell was assigned to a given well if its ClonalBC matched a barcode in that well’s predefined set. Cells whose ClonalBC matched no well’s barcode set were labelled “unassigned” and excluded from the concordance analysis. Only cells present in both the distance matrix and the ClonalBC table, and assigned to a well, were retained. Agreement between the ClonalBC-based well assignment and the lineage tree clade assignment was visualized as a confusion matrix of cell counts for every well-by-clade combination.

### Clonal growth rate analysis

To quantify the relative expansion of individual clones over the in vitro time course, per-clone frequencies and growth rates were computed from ClonalBC counts. Barcode counts were obtained at five parallel-culture time points (Day 17, 24, 31, 38 and 45), from the pre-transplant pool (two Initial captures, combined), and from the in vivo mouse organ samples. Only barcodes of the expected 10 nt length were retained. Within each sample, barcode counts were converted to relative frequencies (percentages of total counts in that sample). Clones were retained if they were detected in the pre-transplant pool, at all five parallel-culture time points, and in at least one in vivo sample, giving 264 clones. For each clone, a per-day growth rate (λ) was estimated by ordinary least-squares regression of log-transformed frequency against time, log(frequency) ∼ day, across the five parallel-culture time points.

### Lineage-tree reconstruction

After the initial validation above, trees were reconstructed one clone at a time, from the intersection of the clone’s cells in the corrected clonal-barcode table with the group edit table. Each tape was treated as a single ordered character rather than six independent sites, and trees were scored by a sequential (prefix) parsimony, a generalization of Camin-Sokal parsimony^52,53^ in which each internal node is labeled by the longest common prefix of its non-missing descendants and the length of a branch is the gain in prefix length from parent to child, summed over tapes. This criterion enforces the irreversibility and the ordering of the recorder and is insensitive to homoplasy at the later sites, which arises when two lineages independently reach a deep site and draw from the same insertion spectrum.

A sequential-aware cell-to-cell distance, the mean per-tape prefix divergence over co-observed tapes, was used to build a neighbor-joining starting tree (Saitou and Nei, Mol Biol Evol, 1987), rooted on an all-unedited outgroup representing the founder. The starting tree was then optimized by nearest-neighbor-interchange hill climbing under the sequential criterion with the founder rooting held fixed. The hill climb was capped at 500 rounds with the global seed and converged well inside that cap in every clone.

*Cross-clone collision screen.* Within each initial clonal edit table, every cell of a clone was screened against the consensus of all other clones of the same mouse. A cell was flagged as a cross-clone clonal-barcode collision when its sequential match to its own clone’s consensus fell below 0.50 while another clone led it by at least 0.20, on at least five co-observed tapes, and flagged cells were pruned from the tree. Pruning is applied once, upstream, and every downstream analysis of that clone inherits the pruned tree, including the transcriptomic within-lineage analyses and the distance-ratio validation. Cell-level quality control was performed uniformly upstream and blind to topology.

*Transition-constrained tree search.* Sequential parsimony is blind to where a cell was sampled. A handful of cells with heavy tape dropout can therefore drift across the lung-liver boundary at essentially no edit cost, and the resulting topology implies several tissue crossings, that is several independent seeding events, as an artifact of that blindness rather than as a claim supported by the edits. A second search stage therefore minimized a penalized objective,

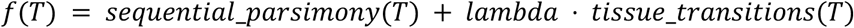

where *tissue_transitions(T)* is the Fitch small-parsimony score^54^ of the discrete sampling-organ label, liver or lung, on *T*, and *lambda* as the regularization parameter. That score is the minimum number of organ changes the topology implies, and is exactly the number of inferred tissue-crossing seeding events. The move set was nearest-neighbor interchange (NNI), evaluated in place and undone so that no tree is copied, together with a guided subtree-prune-and-regraft (SPR) that prunes an organ-pure block whose parent is impure and regrafts it beside the largest clade of the same organ. Each round took the single best improving move, with nearest-neighbor interchange preferred because it is evaluated in place.

The frontier was traced in three passes. First, the same hill climb was run at lambda = 0 with no cap to give an unconstrained reference under the identical move set. Second, a lambda ladder (0, 1, 2, 5, 10, 25, 50, 100, 200, 400, 800, 1600, 3200 insertions per crossing) was walked by continuation, each lambda restarting from the previous lambda’s optimum. Third, because a pure lambda ladder can stall in a local optimum, crossings were peeled off one at a time under a hard cap: the NNI or guided SPR that lowers the crossing count for the smallest parsimony increase was taken, parsimony was then re-optimized under the new cap, and the peel repeated down to k_min. All trees seen in any pass were consolidated into a nested frontier *P(<=k)*, the lowest parsimony reachable while the topology implies at most k crossings, which is monotone by construction so that the difference *P(<=k) – P(<=k+1)* is unambiguously the parsimony the extra crossing buys. The topology reported for each clone is the one adopted at *k = k_min*. Adjacent frontier trees were compared by a tape bootstrap of the paired parsimony difference, δ = *P(<=k) – P(<=k+1)*, with 2,000 resamples and the global seed, following the Kishino-Hasegawa framework^55^.

*Subclonal colony trees.* For the in vitro heritability analysis, full-depth single-cell trees were reconstructed for each monoclonal subclone colony with the Cassiopeia greedy solver^53^ on the colony’s full complement of cells (8 colonies, 313 to 10,506 cells each).

### Recorder-clock calibration and metastasis dating

*Clock model and calibration.* Experimental time was obtained by mapping a node’s cumulative edit depth *d*, the mean number of insertions per recovered tape, to a day through a saturating clock,

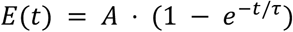

where *A* is the editing asymptote and *τ* the deceleration timescale. The two parameters were calibrated once from independent in vitro time courses of the same recorder line and then held fixed for every dating analysis. Editing data from the bulk-amplicon series from day 17 to day 45 were rescaled onto the single-cell scale by a single multiplicative factor fit jointly with A and tau. The calibration was validated on single-cell data (all harvested at day 45) held out of the fit.

*Dissemination-time estimation.* Each liver metastasis was timed as a unit from its metastatic founder, the most recent common ancestor of the liver cells. The founder’s edit depth was taken as the liver-only consensus, measured per recovered tape to match the clock scale, and converted directly to a dissemination date. Uncertainty was obtained from a tape bootstrap and by varying the consensus threshold over the range 0.6 to 0.8.

*Distance-ratio cross-validation.* As an independent check that shares the clock but not the estimator, the dissemination day was re-estimated from pairwise distances. Let *d* be the sequential parsimony distance between two cells and *E(t)* the saturating clock above, giving edit depth as a function of day *t*. Under that clock the distance between two cells is twice the depth accumulated since their common ancestor, so a pair coalescing at day *t* has expected distance *2[E(T) − E(t)]*, where T = 45 is the harvest day.

For a metastasis founded by a single cell, two liver cells coalesce at the dissemination day *X*, whereas a liver cell and a primary-lung cell coalesce at the clone founder *T_f_*. Their median distances therefore stand in the ratio:

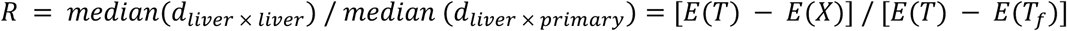

which rearranges to:

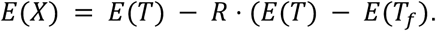

The dissemination day is then recovered by inverting the clock, *X = E^-^*^1^*(E(X))*. The factor of two cancels in *R*, as do technical dropout and homoplasy, which inflate both medians by approximately the same amount. The estimator therefore needs only *R* and two depths: the harvest depth *E(T) = E(45) =* 4.88 edits per recovered tape, and the founder depth *E(T_f_)*, the edit depth at the lung–liver split, taken from each clone’s founder most recent common ancestor. Because X is anchored on that founder rather than on a day-0 root, dropouts that shift all depths together do not bias the estimate. A 95% confidence interval was obtained by a cell bootstrap of 2,000 resamples, and known barcode-collision cells were excluded.

### Migration inference using Metient

To reconstruct the metastatic migration history of each clone, we used Metient (v1.0.0)^23^ to infer the anatomical site of every ancestral node in its phylogeny. Each phylogeny has its leaves labelled with the tissue from which the corresponding cell was sampled, and Metient assigns sites to the unlabelled internal nodes. The analysis was restricted to Mouse 1, the only animal with multi-organ dissemination; Mice 2 and 3 each carry a single focal liver metastasis and are not migration problems. Of the 93 Mouse 1 clones with at least ten recovered cells, 76 were also present in at least two organs. One of these (CGACCAGGAG) lost every cell of its second organ during the cross-clone collision screen and was left with a single anatomical site, which admits no migration history; the remaining 75 clones (5,551 cells) were analysed.

Metient assumes the root of the input tree originated in the primary site, so when the primary is undersampled, a root with few or no primary-site descendants is forced to the primary, spawning a separate migration on every metastatic branch below it. We therefore added a pseudo-root fixed to the primary (LL) above the original root, which counts this departure once and frees the original root to take a metastatic label; when the primary site is adequately sampled and the original root would be labelled primary regardless, the pseudo-root adds no migration. Across the 75 clones this device contributed 43 of 444 total migrations, all of them leaving LL. Because those edges follow from the placement of the pseudo-root rather than from tree structure, proportions reported in the Results are computed over the 401 within-tree migrations. Metient was run in evaluate mode. Alternative migration histories were scored using the pancancer_genetic_uniform_weighting weights, with sample_size = 2048 and num_runs = 3; all other parameters were left at their defaults.

*Calculating tissue transition matrices.* For each clone, we summarized Metient’s multiple solutions as an expected migration graph: each solution’s migration graph (computed using Metient’s migration_graph, whose entries count the migrations between each ordered pair of tumor sites) was weighted by that solution’s probability and summed over solutions, giving the expected number of migrations between each pair of tumor sites. These per-clone graphs were summed across clones and row-normalized, so that each entry gives the probability that a migration departing a given source tissue arrived at a given destination tissue. As Metient returns multiple Pareto-optimal solutions per clone, we took the most probable solution for each clone and counted, for every ordered pair of tissues, the tree edges whose parent node is labelled with the source tissue and whose child is labelled with the destination; this reproduces Metient’s own ‘migration_number’ for all 75 clones. Per-clone matrices were summed across clones and row-normalized, so that each entry gives the probability that a migration departing a given source tissue arrived at a given destination tissue.

*Seeding classifications.* For each clone and each tumor site we counted the migrations arriving at that site in the clone’s most probable solution, and classified the pair as no seeding (0 migrations), single-migration (1) or multi-migration (two or more). Data shown in **Figure 4d** reports, for each site, the percentage of the 75 clones falling in each class. For the primary site, a migration into LL necessarily represents reseeding from a metastatic site rather than initial outgrowth, so that column reflects reseeding of the primary; none of the 155 inferred LN→LL migrations arose in clones lacking sampled primary cells, suggesting that this is not an artefact of the pseudo-root convention.

*Inferring migration histories on A549 data.* Single-cell lineage-tracing data from a Cas9-based lineage recorder were taken from Quinn et al.^6^, using the allele table downloaded from GSM4905334. Of the 100 clones in the allele table, we filtered out 17 clones that failed quality control in the original Quinn et al. study and 4 clones with no cells sampled outside the primary tumor, retaining 79 clones for downstream analysis. For each clone we reconstructed a phylogeny with the VanillaGreedySolver from Cassiopeia^53^ (v2.1.0), using empirical indel priors computed from the allele table and a character matrix built with an allele-representation threshold of 0.9, collapsing mutationless edges. Metient (v1.0.0) was then run in evaluate mode on these 79 phylogenies. Each tree was augmented with a pseudo-root fixed to the primary site (LL) to avoid inflating the inferred migration number, alternative histories were scored using the pancancer_genetic_uniform_weighting weights, and all other Metient parameters were set to their defaults.

### Single-cell lineage and transcriptomic analyses

To analyze the transcriptomic changes during lung-to-liver adaptation, each clone was processed separately, so that an environmental shift cannot be a clone effect. For each of the three liver-metastatic clones that carried both primary-lung and liver cells, a clonally matched triad of single-cell transcriptomes was assembled from the pre-transplant in vitro pool, the lung primary, and the liver metastasis.

*Module scoring.* Each cell was scored with UCell^31^, which is rank-based over the top 1,500 expressed genes per cell and therefore robust to sequencing depth. The candidate set was 66 programs: the 50 MSigDB Hallmark gene sets^26^, the Seurat S-phase and G2M cell-cycle sets^56^, and the pan-cancer cell-state modules of Barkley et al.^25^ after the three neural modules were excluded. Sixty-four were scored.

*Pseudobulk differential expression.* Counts were aggregated by clone and environment and tested with edgeR^57^ under the design ‘∼ clone + env’ with the clone as the replicate, using the quasi-likelihood F test, so that cells are not pseudoreplicated. Only the three genotype-matched triads enter this contrast, giving six pseudobulk profiles and two residual degrees of freedom. Genes were filtered with ‘filterByExpr’ and libraries normalized by trimmed mean of M values. The lung-to-liver contrast tested 14,362 genes, with a common biological coefficient of variation of 0.151, and gave 4,669 genes at 5% false-discovery rate (2,340 up and 2,329 down), of which 1,879 also changed at least two-fold (458 up and 1,421 down).

*Gene-set enrichment.* fgsea^58^ was run on the full ranked differential-expression list against the Hallmark, Barkley pan-cancer cell-state, and MSigDB C6 oncogenic collections, ranking genes by the signed log P value. Because the pooled enrichment answers whether a program is enriched in one averaged ranking rather than whether the shift reproduces across clonal genotypes, each program was additionally resolved per clone as the liver-minus-lung difference in mean UCell score, with a 95% confidence interval from the per-cell scores and an across-clone paired t-test at n = 3. The per-clone UCell difference is displayed rather than a per-clone enrichment score because a within-clone ranking has no environment replicate and fgsea returns no score for the strongest proliferation sets in the smallest clone.

*Transcription-factor activity.* Per-cell transcription-factor activity was inferred with decoupleR^27^ using the univariate linear model method over the CollecTRI regulatory network^28^, and pseudobulked per clone and environment. The network covers 1,186 transcription factors, of which 697 had at least five target genes expressed in this dataset and were scored; the remaining 489 were dropped for insufficient targets. Regulators were ranked by the effect size that is consistent across all three clones.

*Lineage-informed co-expression modules.* Hotspot^29^ was run with the edit-scaled single-cell lineage tree as the cell-to-cell metric and the danb model, per clone, with the mitochondrial and ribosomal cell-quality signal removed and module scores residualized on mitochondrial fraction and log total counts. Consensus modules across clones were formed by taking the recurrent lineage-autocorrelated genes; each clone’s local co-expression matrix was averaged, and the consensus matrix was clustered. Because a metastasis founded by a single cell is itself a monophyletic clade, the deepest split on a whole-tumor tree is the lung-liver boundary and the whole-tumor Hotspot signal coincides with the culture-to-liver transition; it therefore corroborates the gene sets recovered by differential expression and fitness but does not on its own establish heritability. Hotspot was consequently also run within the lung-only and liver-only subtrees, where lineage and environment are decoupled, and on the in vitro monoclonal colonies.

*Phylogenetic fitness.* For each of the three metastatic clones, a per-cell phylogenetic fitness was estimated from the shape of its edit-scaled single-cell lineage tree using the local branching index^30^, computed with a neighborhood scale tau = 0.0625 times the maximum patristic distance and normalized to a maximum of 1, so that densely and recently branching lineages score highest. To identify the transcriptional programs associated with fitness, each gene’s expression was correlated with per-cell local branching index (Spearman) across the cells of each clone, restricting to genes detected in at least 20% of that clone’s cells. Because the index covaries with cell quality and cell-cycle activity, these were recomputed as partial correlations, rank-residualizing both expression and fitness on log total counts, mitochondrial fraction, and S-phase and G2M cell-cycle scores^56^ before correlating, per clone. A gene was called fitness-associated if the partial correlation had a consistent sign across all three clones and a mean value above 0.2, giving 212 genes of the 10,179 tested in every clone (**Supplementary Table S5**). Where a compact score was needed rather than the whole set, the top thirty genes of that ordering were used as the InvasionFitness signature, which is what the Mouse 1 clone table is scored against; it is a strict subset of the fitness-associated genes, and **Supplementary Table S5** flags the two sets separately because they are not interchangeable. Controlling for these covariates collapsed the proliferation and OXPHOS associations to near zero in every clone while the partial-EMT, hypoxia, and glycolysis associations survived, so the fitness-correlated lineages are not simply the most proliferative. For the program-level comparison in **Figure 5h**, each program’s fitness association was quantified as the enrichment of its gene set among the fitness-associated genes, expressed as –log10 P from a hypergeometric test against the expressed transcriptome (approximately 10,200 genes). This value is the y-axis of **Figure 5h** and the fitness column of the **Supplementary Table S6**; the enrichment backgrounds in that matrix are approximate and the matrix is used to display the pattern within each metric, not to compare values across metrics.

*In vitro lineage heritability.* The environment-decoupled test of heritability was run on the monoclonal subclone colonies, a single homogeneous environment in which lineage is decoupled from tissue and organ-niche gradients. The same lineage-informed module analysis was applied to 8 full-depth colony trees built using Cassiopeia^53^ (313 to 10,506 cells each). Each colony showed hundreds to thousands of lineage-autocorrelated genes, with the number scaling with colony size, and the same partial-EMT/adhesion and glycolysis modules recurred in every colony with at least one module; only the smallest colony was power-limited. Genes reproducibly heritable across colonies were defined as those significant at Hotspot false-discovery rate below 0.05 in at least 75% of the colonies that tested them, requiring at least four colonies tested, giving a 396-gene set. Heritability was estimated independently by PATH^32^ (phylogenetic autocorrelation over all pairs of tips, inverse tree distance weighting), applying the same fractional rule, giving 611 genes. The two estimators agree over the whole range (Spearman rho = 0.86 across the 10,078 genes scored by both), and their intersection of 349 genes is taken as the heritable set (**Supplementary Table S7**).

*Scoring clones from the pre-transplant for gene programs.* Each cell from the Pre-TX pool was assigned to its clonal barcode by highest-UMI resolution and depth-normalized, yielding 3,208 barcoded clones over 39,082 cells. Each cell was scored with UCell for six signatures: (i) the pre-existing state, the 349 two-method heritable genes defined above (348 of which are present in the expression matrix); (ii) glycolysis (HALLMARK_GLYCOLYSIS); (iii) hypoxia (HALLMARK_HYPOXIA); (iv) partial EMT from Barkley et al.^25^; and, as specificity controls, two programs that are strongly upregulated in the liver but neither fitness-associated nor heritable, namely (vi) proliferation (MKI67, TOP2A, MCM2, MCM6, PCNA, CCNB1, CDK1, TYMS, MYC) and (vii) OXPHOS (HALLMARK_OXIDATIVE_PHOSPHORYLATION). Both controls behave as intended: OXPHOS and hypoxia are null across every outcome, and proliferation reaches nominal significance in Mouse 1 and Mouse 2 (1.25 and 1.20, both P about 0.05, with confidence intervals bounded at one) but not Mouse 3 and not for dissemination, so it is not reproducible across mice rather than null (**Supplementary Figure 8c**). Each clone was summarized by the mean of each score across its pre-transplant cells, restricted to the 1,229 clones with at least 10 cells (31,838 cells) for a stable per-clone estimate, together with its pool abundance, the number of pre-transplant cells recovered for that clone.

*Modeling primary colonization and dissemination.* Because the same pool was transplanted into all three mice, several recovery outcomes were scored per clone: colonization of the primary lung in Mouse 1, Mouse 2, and Mouse 3 separately; metastasis pooled across the three mice, defined as recovery at any secondary site; and, for the single-mouse multivariable model, recovery anywhere in the polyclonal Mouse 1 cohort. For each signature and outcome a logistic regression was fit, outcome ∼ log(pool abundance) + scale(signature), and the odds ratio per standard deviation of state is reported as the exponentiated scaled-signature coefficient with its 95% Wald interval and P value, alongside a likelihood-ratio test against the abundance-only model. Conditioning on log pool abundance controls for the sampling bottleneck, since more abundant clones are more likely to be recovered irrespective of state. For multivariate modeling, glycolysis, the pre-existing heritable set and partial EMT were fitted jointly on each outcome (outcome ∼ log(pool abundance) + glycolysis + pre-existing state + partial EMT; **Supplementary Figure 8d**).

*Per-gene competence scan.* To test the competence of individual genes, each pre-transplant clone with at least 10 barcoded cells was reduced to a single pseudobulk profile by summing counts across its cells, giving 1,229 clones over 31,838 cells, and counts were converted to log2 counts per million. Genes detected in at least 5% of clones and non-constant across them were retained, leaving 16,221. Two clone-level outcomes were defined: establishment, recovery at any site in any mouse against recovery nowhere, and dissemination, recovery at a secondary site against recovery only in a lung, the latter restricted to established clones so that it does not re-measure the first. Significance was assessed against a background generated by reshuffling the outcome labels across clones (200 rounds), which holds the expression matrix and its correlation structure fixed and destroys only the clone-to-outcome link; the 5% false-discovery threshold against that background is *P* = 6.6 x 10^-5^.

### Low-TAPE/TargetBC recovery in vivo cell population

Twelve tumor samples were analyzed together: the six Mouse 1 organs and the lung and liver of Mouse 2 and Mouse 3. Each cell was labeled recovered or non-recovered by whether its cell barcode appeared in that sample’s clonal-barcode recovery table. Each cell was scored with UCell (maxRank 1,500) for a cell-cycle signature (the union of the Seurat S and G2M gene sets), a curated dormancy signature (*NR2F1, BHLHE41, THBS1, TGFB2, DKK1, CDKN1A, CDKN1B, SOX9, AXL, TXNIP, DDIT4, ZFP36, KLF6*), a curated G0-quiescence signature (*CDKN1A, CDKN1B, RB1, FOXO3, GADD45A/B, BTG1/2, KLF4, KLF6, ZFP36, CEBPB, TXNIP, DDIT4, NR2F1*), the pan-cancer pEMT module, and the cytoplasmic ribosomal protein genes (*RPS* and *RPL*). Cell-cycle phase was assigned with Seurat “CellCycleScoring” on the same S and G2M sets. To distinguish a quiescent state from a low-RNA artifact, recovered cells were downsampled by binomial UMI thinning to the non-recovered median total count of their own sample (mouse and tissue, the same unit used for the pseudobulk contrast) and rescored, and a logistic regression of recovery on scaled log total counts, cell-cycle score, and dormancy score with tissue as a covariate was fit. Recovered cells carry a median 27,854 UMIs against 2,578 in non-recovered cells, a 10.8-fold difference, and recovery is dominated by transcriptional output (odds ratio 8.6 per standard deviation of log total counts, 95% confidence interval 8.3 to 9.0). Pseudobulk differential expression between recovered and non-recovered cells was computed with edgeR (quasi-likelihood F test on a ‘∼ sample + recovery’ design, so each sample serves as its own paired control) in a combined model and in lung-only, liver-only, and depth-matched variants. A sample enters the pseudobulk only if it contributes at least twenty cells on both sides, which excludes the 44-cell Mouse 1 intraperitoneal sample (16 recovered) and leaves eleven of the twelve. The lung and liver recovery signatures were compared by the Spearman correlation of their gene-wise fold changes, and reported as **Supplementary Figure 3** and the accompanying **Supplementary Note 1**.

### Statistical analysis

Unless otherwise stated, the statistical unit for the transcriptomic contrasts is the clone, not the cell, and pseudobulk aggregation with the clone as replicate was used to avoid pseudoreplication. Gene-level and gene-set tests report Benjamini-Hochberg false-discovery rates. Logistic-regression coefficients are reported as odds ratios per standard deviation of the predictor with 95% Wald intervals and two-sided *P* values, alongside likelihood-ratio tests against the nested model. Correlations are Spearman unless stated otherwise, and partial correlations were computed by rank residualization on the stated covariates. Confidence intervals on tree-derived quantities were obtained by bootstrap, resampling tapes for parsimony and clock quantities and cells for the distance-ratio estimator, with 2,000 resamples. Comparisons involving three clones or three mice are reported with the *n* stated in the relevant legend and are treated as reproducibility rather than as powered tests. P values from single uncorrected outcomes are identified as such. Randomized steps used fixed seeds as specified in the shared analysis script, and the distance, sequential-parsimony, and frontier steps are deterministic. The clonal-barcode collapse breaks ties alphabetically and the insertion-color palette in the tree panels is generated from the alphabetically sorted set of unique insertions. Analyses used Python 3.10 (Cassiopeia^53^ 2.1.0, numba 0.60, ete3 3.1.3, numpy 2.4, scipy 1.17, pandas 2.3) for the lineage reconstruction, a separate Python environment for the transcriptomic analysis (scanpy 1.10.3, anndata 0.10.9, decoupleR^27^ 1.8.0, hotspotsc^29^ 1.1.3, numpy 1.26.4), and R 4.3.3 with Seurat^51^ 5.2.1, UCell^31^ 2.6.2, msigdbr^26^ 25.1.0, edgeR^57^ 4.0.16, fgsea^58^ 1.28.0, SingleCellExperiment^59^ 1.24.0, ape 5.8.1^60^, phangorn 2.12.1^61^, and ggtree 3.10.1^62^.

### AI disclosure statement

We disclose that data exploration, data analysis, and manuscript writing were supported by AI-based tools. The authors take full responsibility for the data, code, analyses, conclusions, and writing.

## DATA AVAILABILITY

Raw sequencing data have been deposited to SRA (BioProject PRJNA1495924), and associated processed data have been deposited in the Gene Expression Omnibus (GEO) under accession number GSE339406 (reviewer token: gringoeqvdafzan). Plasmids generated in this study have been deposited with Addgene and are available under the following plasmid numbers: PB-EF1a-loxP-HygR-3xPolyA-loxP-PE2max-P2A-mClover3, Addgene #261083; Lenti-EF1a-Puro-P2A-Luciferase-WPRE, Addgene #261084; PB-U6-pegRNA-BbsI-BbsI-EF1a-mRFP-TAPE-BsmBI-BsmBI, Addgene #261085; pLenti-CAG-Cre-T2A-mRuby3-WPRE, Addgene #261086.

## CODE AVAILABILITY

Metient-related scripts are available at GitHub (https://github.com/divyakoyy/dna_typewriter_metastatic_histories), and all other analysis scripts are available at GitHub (https://github.com/choi-junhong/lung-xenograft-lineage), archived at Zenodo (https://doi.org/10.5281/zenodo.21875723).

## SUPPLEMENTARY INFORMATION

**Supplementary Table S1. Nucleic acid sequences used in the study.** Sequences of the prime-editor and pegRNA constructs, the 6xTAPE array and its TargetBC region, the static clonal barcode, and the primers used for targeted amplification of the recorder and barcode libraries.

**Supplementary Table S2. Clonal barcodes after assignment.** The corrected clonal barcode called for each cell, with its sample of origin, after Hamming-distance-one collapse of the raw barcode calls. This is the assignment on which every clonal analysis in the paper rests.

**Supplementary Table S3. Lung-to-liver differential expression across the three clonally matched metastases.** Gene-level differential expression between the liver metastases and the clonally matched lung primaries, computed with edgeR using the clone as the replicate unit. Reports the log fold change, *P* value and false-discovery rate for every gene tested, with positive values indicating higher expression in the liver. The 4,669 genes at 5% false-discovery rate and the 1,879 changing at least two-fold (458 up, 1,421 down) quoted in the text are subsets of this table.

**Supplementary Table S4. Composition of the five named transcriptional programs.** Each program used in Figures 5 and 6 mapped to its constituent MSigDB Hallmark sets with their normalized enrichment scores in the liver-versus-lung contrast, and how many of the sets tested reached significance; its pan-cancer cell-state module (Barkley et al.) with normalized enrichment score and adjusted P value; and its associated CollecTRI regulons. This table defines what each program name refers to throughout the paper.

**Supplementary Table S5. Genes associated with phylogenetic fitness within the liver metastases.** Per-clone and mean Spearman correlations between gene expression and the per-cell local branching index, both naive and partial, the latter rank-residualizing expression and fitness on log total counts, mitochondrial fraction and S-phase and G2M cell-cycle scores. Covers every gene detected in at least 20% of the cells of all three dated clones. Two nested sets are flagged separately and are not interchangeable: “fitness_associated” marks the 212 genes whose partial correlation has a consistent sign in all three clones and a mean above 0.2, which is the extracellular-matrix, adhesion and invasion set named in the text; “invasion_signature” marks the compact top thirty of that ordering, which is what the Mouse 1 clone table is scored against.

**Supplementary Table S6. Each transcriptional program placed on three independent axes.** For the five programs: liver-adaptation strength, as the normalized enrichment score in the liver-versus-lung contrast; in vitro lineage heritability, as the enrichment of the program’s genes among the heritable set, expressed as *-log10 P* from a hypergeometric test; and phylogenetic fitness association, as the mean partial Spearman correlation with the local branching index. The three metrics use different enrichment backgrounds, so values are comparable within a column but not across columns.

**Supplementary Table S7. Recurrently lineage-heritable genes in the pre-transplant colonies.** The 349 genes are called lineage-heritable in the monoclonal pre-transplant colonies by both methods, which is the set used as the pre-existing signature throughout Figure 6. For each gene: the number of colonies in which it was tested and the number in which it was significant, under Hotspot and under PATH separately; the fractional-recurrence call from each method; the program it belongs to; its association with establishment; and flags marking membership of the two-method intersection. The recurrence rule is: significance in at least 75% of the colonies that tested a gene, tested in at least four.

**Supplementary Table S8. Per-clone pre-transplant scores and competence-prediction model outputs.** This includes a contents sheet and three data sheets: *Per-clone scores*: one row for each of the 1,229 pre-transplant clones, giving its abundance in the pool, its score for every program and for the heritable set, and the outcomes it was scored against; this is the table every Figure 6d and 6e model is fitted on. *Mouse 1 clone table*: 305 clones with dissemination breadth, invasion score and phylogenetic fitness across the Mouse 1 organs. *Model fits*: 120 fitted models, giving the odds ratio per standard deviation with its 95% confidence interval and P value for each outcome and signature, with the first column naming the scope (single-mouse multivariable, multi-mouse abundance-adjusted, multi-mouse odds ratios). Outcomes follow the nested contrast structure throughout, with establishment scored as colonization anywhere against never, and dissemination as reaching a secondary site against lung-only among the clones that established.

**Supplementary Note 1. Low-TAPE/TargetBC recovery in vivo cell population**.

All three mice and both the primary lung and the liver metastases were analyzed together (68,691 single cells from twelve samples; **Methods**). Low recovery was pervasive: 77% of transcriptomically defined tumor cells yielded no clonal barcode. The fraction varied by site along a reproducible gradient (**Supplementary Figure 3a**), highest in the lung primary (86 to 95%) and the peripheral thoracic sites (lymph node, thoracic mass, and intraperitoneal nodes, 64 to 67%) and lowest in the liver metastasis (39% in Mouse 2, 64% in Mouse 3).

Relative to recovered cells they had roughly 10-fold lower total RNA, about 5-fold fewer detected genes, markedly lower ribosomal and translational gene expression, and a lower cycling fraction (34% versus 46% in S or G2M) (**Supplementary Figure 3b,c**). Because recovery of the clonal barcode and DNA Tape is an amplicon measurement that scales with transcriptional output, recovery is itself largely a readout of that output: in a logistic model the odds of recovery rose roughly eight-fold per standard deviation of total RNA (**Supplementary Figure 3d**).

To characterize the state without presupposing a marker set, we applied the same module-enrichment approach used for the liver adaptation, testing paired pseudobulk differential expression (recovered versus non-recovered within each sample) against the Hallmark and pan-cancer cell-state collections (**Supplementary Figure 3e**; **Methods**). The dominant and most consistent feature was a coordinate shutdown of proliferation and biosynthesis in the low-recovery cells: *MYC* targets (normalized enrichment score or NES = –3.0), E2F targets (–2.2), the G2M checkpoint (–1.5), DNA repair (–1.6) and OXPHOS (–1.7) were all depleted, and the pan-cancer cell-cycle module was the most depleted module in every contrast (NES = –3.0 depth-matched, –2.9 in the lung, –3.5 in the liver). A published quiescent-versus-dividing signature was independently enriched (NES = 1.8, FDR = 1 × 10^-4^).

